# Unravelling biomolecular and community grammars of RNA granules via machine learning

**DOI:** 10.1101/2024.04.06.588388

**Authors:** Zhan Ban, Yan Yan, Kenneth A. Dawson

## Abstract

RNA granules are dynamic compartments within cells that play a crucial role in posttranscriptional regulation of gene expression. They are associated with a variety of human neurodegenerative diseases. While RNA granules play vital roles in cellular functions, the comprehension of their assembly has remained elusive.

In this study, we employed robust machine learning models combining residue content and physicochemical features to accurately identify potential RNA granule (*i.e.,* stress granule and P-body) proteome within the human proteome. Our models achieved good performance with high areas under the receiver operating characteristic curve of up to 0.88, outperforming previous liquid-liquid phase separation models. Intriguingly, the predicted RNA granule proteome reveals a significant enrichment in biological functions and domains associated with RNA granule-related processes, mirroring findings from observed high-confidence RNA granule protein datasets. Furthermore, our analysis unveils critical physicochemical attributes, notably hydrophobicity, influencing the formation of RNA granules.

Using the constructed model, we uncovered the central roles of RNA granule proteins with high propensities within the comprehensive RNA granule protein-protein interaction (PPI) network and their commonality in diverse RNA granules. Furthermore, we identified prominent clusters with dense PPIs, significantly contributing to critical biological processes within diverse RNA granules, including translation, mRNA decay, rRNA processing, and mRNA splicing. This analysis proposes a hypothesis: dense PPI clusters are integral functional subunits, constituting relatively stable ‘cores’ within diverse RNA granules.

In conclusion, this study provides a comprehensive molecular and community-based foundation for understanding the importance of PPIs in the stability of RNA granule formation and functionality. This analysis contributes to a deeper and more comprehensive understanding of the intricate nature of RNA granules and opens avenues for future research and therapeutic interventions targeting RNA granule- related diseases.

## Main

Compartmentalization of molecular constituents is a fundamental process for cells to maintain and control numerous biochemical processes simultaneously^1^. Apart from the membrane-bound organelles, such as the endoplasmic reticulum (ER), mitochondria, and nucleus, there are biomolecular organelles without membranes, which are found ubiquitously, such as RNA granules. They are formed via liquid-liquid phase separation (LLPS)^2^ within cells, which is driven by the attractive interactions among components according to the general glass paradigm in soft matter and colloid science^3^.

RNA granules are heterogeneous aggregates that comprise various RNA and RNA binding proteins (RBPs), ribosomal subunits, decay enzymes and translation factors^4^. They play a crucial role in maintaining a wide range of functions, such as post-transcriptional regulation of gene expression^4^ and stabilization of their RNA cargo^4^. Moreover, increasing evidence suggests that dysregulation of RNA granules is associated with various human diseases, including frontotemporal lobar degeneration and amyotrophic lateral sclerosis^5,6^.

Despite significant strides in RNA granule research, understanding their formation and functionality remains challenging due to the dynamic and stress-dependent nature of their components^7^. RNA granule components determine their functions^8–10^. Their highly dynamic components may enable RNA granules to respond to stress specifically. In addition to the crucial roles of RNA granules in fundamental biological processes^4^, RNA granules seem to exhibit a subtle but bold strategy to balance between stability and specificity of their formation and functionality in complex cellular environments, without the help of membranes.

To advance our understanding, one critical area of investigation focuses on identifying RNA granule proteome and elucidating the underlying biomolecular patterns governing their formation and functionality. Recent resource-costing and timing-consuming methods, such as co-localization with established RNA granule markers and catalog construction^1^, limit the identification of RNA granule components. However, recent RNA granule studies^1^ tend to gain theoretical guidance from machine learning models on LLPS^11–13^, one of the driving forces in RNA granule formation, but lacking specificity and reliability in identifying the composition of RNA granules. Therefore, despite the advancements made in RNA granule studies and LLPS model building, there is still a significant knowledge gap concerning a deeper understanding of the underlying biomolecular grammars in specific RNA granules.

Recent studies, such as those by Jain et al.^14^ and Markmiller et al.^7^, shed light on relatively stable ‘core’ substructures and dense protein-protein interaction (PPI) networks within stress granules (SGs). For example, Jain and colleagues reported that relatively stable core substructures were observed surrounded by a phase-separated shell^14^. The cores may maintain the stability of RNA granule functionality via dense PPIs, while the dynamic shell-like phase-separated structure contributes to their assembly, disassembly and transitions^14^. Furthermore, in even unstressed cells, a highly dense and pre-existing SG proteome PPI network was also observed^7^. Some essential RNA granule proteins with important biological functions (*e.g.,* UPF1, AGO2 and HNRNPU) were found in the PPI network with high centralities^7^. Therefore, dense PPIs may play an important role in RNA granule functionality. However, the previous individual experiments could only provide narrow perspectives on the highly dynamic and heterogeneous RNA granule systems.

This study introduces a novel approach that integrates sequence-based protein features with observed high-confidence RNA granule proteins to accurately identify the RNA granule proteome within the human proteome. Despite the limited number of high-confidence proteins available in the RNA granule database (*e.g.*, 280 tier 1 proteins for the SG)^1^, our approach yields RNA granule models with good performance, achieving the areas under the receiver operating characteristic (ROC) curve (AUCs) of up to 0.88, outperforming previous liquid-liquid phase separation (LLPS) models. Significantly, the predicted RNA granule proteome could cover most of the collected heterogeneous RNA granule proteomes published from diverse experiments (77%). Meanwhile, the predicted RNA granule proteome is significantly enriched in the crucial biological functional terms in relation to RNA granule’s functions, such as RNA transport and spliceosome pathways, thereby aligning well with the enrichment results of observed high-confidence RNA granule proteins. Overall, our study provides a molecularly principled foundation (*e.g.,* hydrophobicity as a key driver) for a deeper understanding of RNA granule formation behaviour, including LLPS.

In addition, we address this knowledge gap by applying an up-to-date RNA granule model to construct an overall identified RNA granule proteome PPI network. Combining predicted biomolecular propensities with graph theory, we delineate the distinct biomolecular and community grammars in RNA granule formation and functionality. Our analysis reveals the central role of crucial RNA granule components in the dense PPI network, contributing to the fundamental functions of RNA granules. Moreover, we identify dense RNA granule clusters intricately linked with fundamental biological functions, offering comprehensive insights into community-level processes such as translation, mRNA decay, rRNA processing, and mRNA splicing. This research enhances our understanding of RNA granules and offers a novel workflow for evaluating complex biological systems through the lens of an accurate machine learning model.

## Results

### Differential Features of RNA Granule Proteins with the Human Proteome

For the assessment of protein sequences regarding their potential to form RNA granules (*i.e.,* SG, P-body (PB), and PB or SG (PBSG)), we utilized an essential RNA granule database, the RNAgranuleDB^1^, that includes human SG and PB proteins as positive samples in our models (see **Table S1**). Negative protein candidate databases were randomly generated from the human proteome. Crucially, specific criteria (ranging from tier 1 to tier 3 or tier 4) are in place to classify RNA granule proteins. These criteria consider diverse evidential factors, including strong co-localization with established SG, PB markers (tier 1) to weak predicted RBP propensity (tier 4) as detailed in the original literature^1^. In our analysis, classification models were developed using high-confidence RNA granule proteins, such as tier 1 proteins of SG (N=280 tier 1 SG proteins + 280 negative proteins) and PBSG (N=473 tier 1 PBSG proteins + 473 negative proteins), or a combination of tier 1 and tier 2 proteins of PB (N=198 tier 1&2 PB proteins + 198 negative proteins). To evaluate the specificity of RNA granule components, we also introduced the collected RBP proteome (N=6163) from the RBPbase database.

Building upon prior machine learning models addressing protein condensate propensity^11^ and PPI models^15,16^, this study employed three protein feature representation methods (**Fig. 1**): physicochemical features, aa content and k-mer content. The physicochemical features (N=19) of proteins, estimated from protein sequences, encompassed fundamental properties such as length, molecular weight (mw, daltons), isoelectric point (IEP), hydrophobicity (or hydrophilicity), aromatic fraction, cation fraction, and fractions of protein secondary structures (*i.e.,* alpha-helix, beta-turn, beta-sheet). Additionally, metrics related to protein sequence entropy (*i.e.,* Shannon entropy) and low complexity regions (LCRs, LCR fraction, LCR scores, and lowest complexity score) were included.

**Fig. 1.**
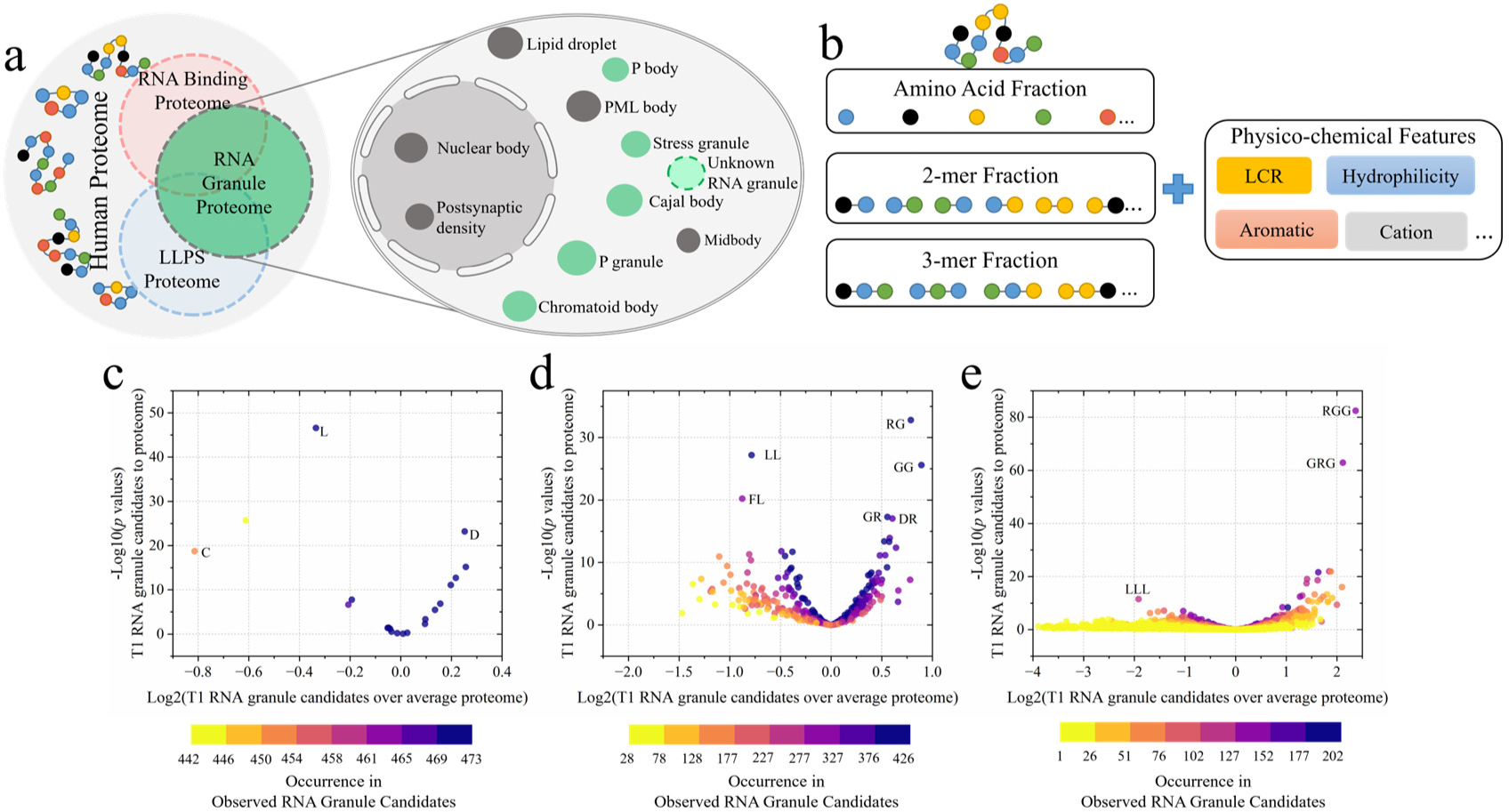
Workflows of RNA granule model construction. To distinguish RNA granule components from human proteome and RBP proteome via machine learning models (**a**), we employed three methods to extract essential protein features (**b**). These included physicochemical features (N=19 in **Fig. S1**) such as LCR fraction, gravy value (*i.e.,* hydrophobicity or hydrophilicity), aromatic fraction and cation fraction; the amino acid (aa) contents (*i.e.,* 1-mer fraction) of the whole protein sequences (N=20 in **c**) and the aa contents in the LCR sequences (N=20) and the k-mer features (*i.e.,* 2-mers: N = 50 in **d**; 3-mers: N=50, corresponding to their *p* values<0.001 compared observed protein candidates with human proteome and occurrence in observed protein candidates, in **e**) that correspond to substrings in protein sequences. Then, we also compared the aa contents, 2-mer contents and 3-mer contents of observed RNA granule protein candidates with collected RBPs in **Fig. S2&S3**. We applied the one-way ANOVA to calculate the *p* values. LCR: low complexity region. T1: tier 1. C: cysteine. L: leucine. D: aspartate. FL: phenylalanine-leucine. LL: leucine-leucine. DR: aspartate-arginine. GR: glycine-arginine. GG: glycine-glycine. LLL: leucine-leucine. GRG: glycine- arginine-glycine. RGG: arginine-glycine-glycine.

The distinct distributions of selected fundamental physicochemical properties are evident between observed RNA granule protein candidates and the human proteome (excluded the collected RNA granule candidates, as illustrated in **Fig. S1**). The high- confidence RNA granule proteins tend to be more hydrophobic (*i.e.,* less gravy values), larger (*i.e.,* higher mws) and more disordered (*i.e.,* with more LCR fractions) compared with human proteome (*p* values < 0.001).

However, compared with previous two classic LLPS models, the DeePhase^12^ and PSAP^11^, our observations (*i.e.,* rich large and hydrophobic proteins in specific RNA granules) are familiar (*i.e.,* rich RGG 3-mers and more disordered regions) and distinct (*i.e.,* rich small and hydrophilic aas) with their observations and conclusions on different features. Crucial elements such as prion-like domains, RGG motifs, and RG repeats within RNA granules collectively contribute to facilitating LLPS processes via a combination of weak multivalent interactions, including hydrophobic interactions^17^. Moreover, there are lots of studies revealed that the presence of IDRs with large and highly hydrophobic residues, plays an important role in LLPS formation and RNA binding processes of some important key RNA granule proteins (*e.g.,* FUS^17,18^ and DDX6^19^). The similarity indicates LLPS proteins and RNA granule proteins could undergo phase separation, a process where a homogeneous solution separates into two distinct phases. While the disparities in multiple physicochemical properties may suggest the unique biomolecular grammars in RNA granule formation to general biomolecular condensates driven by LLPS.

K-mers, substrings of length k within biological sequences, are commonly used in sequence analysis, including PPIs^15,20^, and the evaluation of RNA-protein interactions^21,22^. In this study, we extracted distinct (*p* values < 0.001) and abundant (according to the top k-mers with highest abundance among RNA granule protein candidates) 2-mer (N=50) and 3-mer (N=50) features, representing fractions of each k-mer in entire sequences, to characterize each protein. Before we developed our models for RNA granules, most research in this area used LLPS models and RBP databases for theoretical guidance^1^. To further evaluate the specific biomolecular grammars of RNA granules, we compared their k-mer features with those of the human proteome (**Fig. 1c-e**) and collected RBP proteome (N=6163, **Fig. S2&S3** and **Table S1**). We observe a substantial and statistically significant abundance of specific residues, such as RG, GG, RGG, and GRG, in tier 1 RNA granule protein candidates when compared to the human proteome and RBP proteome. Notably, the enriched contents of RGG/RG motifs, including RG, GG, and RGG with GRG, are prevalent in IDRs of proteins, such as FUS, playing a vital role in LLPS processes and facilitating weak multivalent interactions with RNA or DNA^17^. Moreover, it’s worth highlighting that over 100 human proteins feature two or more RGG motifs, each separated by fewer than 5 residues^23^. The significance of multiple RGG/RG motif repeats at the molecular level becomes evident when considering their crucial roles in facilitating high-affinity interactions between proteins and RNA structures^17^. This observation illuminates the potential of our feature representation methods in machine learning to capture the existence of multivalent and dynamic interactions at the molecular level (**Fig. S4&S5**).

In addition to the abundance of hydrophobic RGG (with gravy value of -1.77) in high-confidence RNA granule protein candidates, the contents of hydrophilic k-mers (*i.e.,* LLL with gravy value of 3.8 in **Fig. 1e** and LL with gravy value of 3.8 in **Fig. 1d**) with hydrophilic residues (*i.e.,* L with gravy value of 3.8 in **Fig. 1c**) tend to be significantly (*p* values < 0.001) less in tier 1 RNA granule protein candidates compared to human proteome. The observations align with observed crucial roles of hydrophobicity in RNA granule formation^24,25^.

However, despite extensive analysis, we did not identify a significantly different distribution of k-mers associated with RGG/RG motifs across overall RNA granule protein candidates (tier 1 to tier 4) and RBPs. This suggests that high-confidence RNA granule proteins (tier 1), often found in the cores of RNA granules, may exhibit unique characteristics compared with other RNA granule components (**Fig. S5**), consistent with the observed significance of stable ’cores’ in SGs’ formation and functionality^14^.

### Robust Machine Learning Models for Distinguishing RNA Granule Proteins from Human Proteome

With selected physicochemical properties and contents of aa with k-mers of proteins, we performed the RNA granule model to distinguish RNA granule proteins (*i.e.,* SG, PB and PBSG proteins) from the human proteome, trained by high-confidence RNA granule protein candidates (*i.e.,* tier 1 for SG and PBSG, tier 1&2 for PB) from the RNAgranuleDB and randomly selected negative proteins from human proteome.

Initially, we evaluated the robustness of the constructed RNA granule models as shown in **Fig. 2a, S6-S8**. We applied the ten-fold cross validation method^28^ to measure models’ robustness. We divided the overall dataset (*i.e.,* training and testing sets) into ten equally sized subsets (*i.e.,* folds). Each fold gets a turn to serve as the validation set, while the remaining nine folds form the training set. This process is iterated ten times, ensuring each fold is used for validation once. This method provides a comprehensive evaluation of the model’s performance across different data samples. According to the ten-fold cross validation accuracy, we found our RNA granule models could achieve robust and stable performance on testing sets (with high AUC values 0.88 ± 0.03 for SG, 0.86 ± 0.08 for PB and 0.87 ± 0.05 for PBSG; with high area under precision-Recall curve (PR AUC) values 0.87 ± 0.05 for SG, 0.86 ± 0.10 for PB and 0.86 ± 0.05 for PBSG in **Fig. S6&S7**) and the unlikely-LLPS Protein Data Bank (PDB) proteins (with high accuracy up to 0.87 ± 0.02 in **Fig. S7**). In contrast to conventional and proteome- applied LLPS propensity prediction models (*e.g.,* PSAP’s AUC of 0.88 with PR AUC of 0.25, and PScore’s AUC of 0.84 with PR AUC of 0.11)^11^, our RNA granule models are performing similarly in terms of overall accuracy, but has improved precision and recall, reducing false positives (*i.e.,* false RNA granule proteins predicted) and detecting more true positives (*i.e.,* true RNA granule proteins predicted). It indicates that the RNA granule models demonstrated superior performance.

**Fig. 2.**
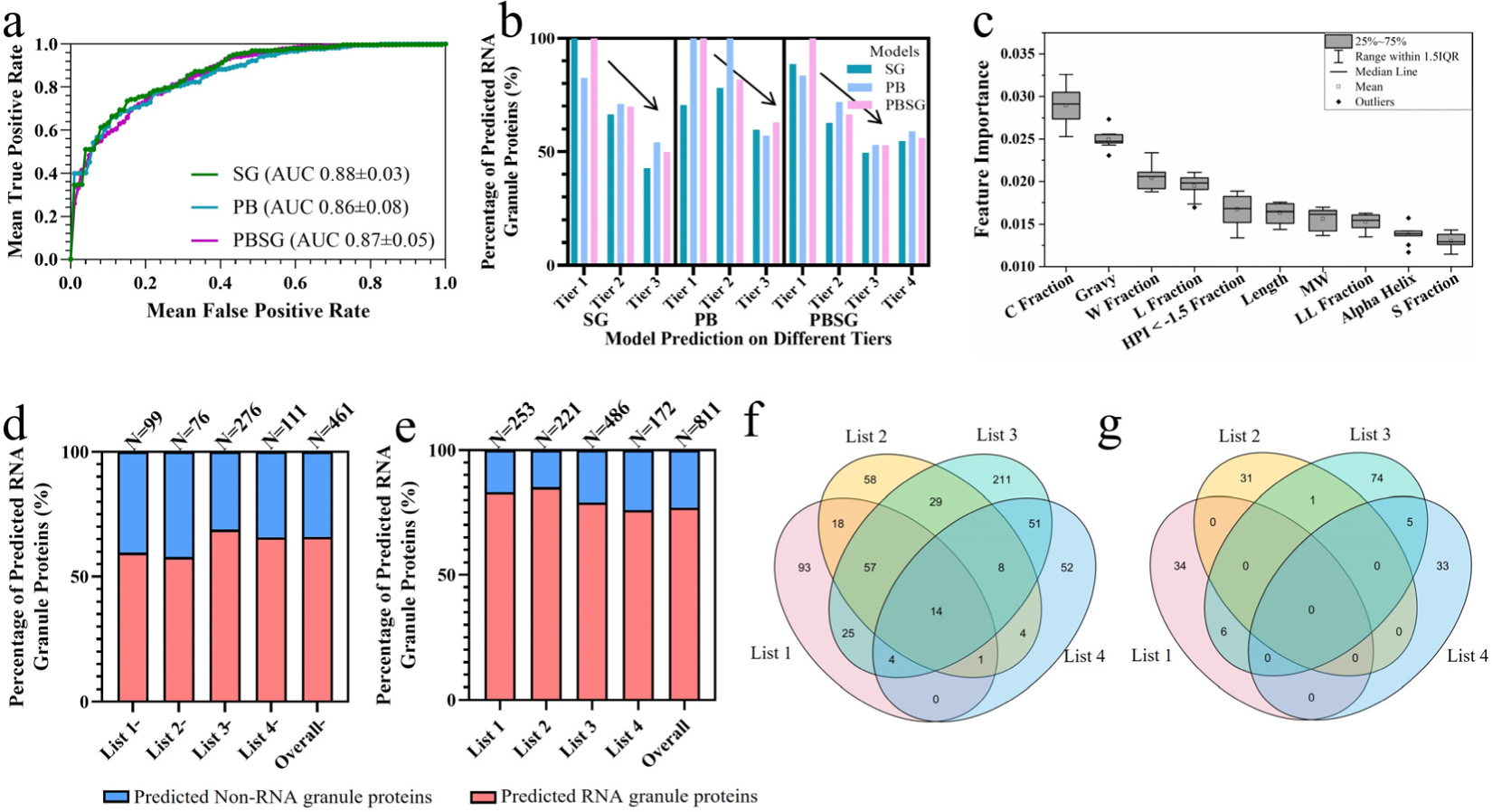
Robustness analysis of RNA granule protein models. (**a**) The analysis applied the ten-fold cross validation method to estimate the model prediction performance, measured by the AUC values (shown in Mean±SEM, N=10) for SG, PB and PBSG proteins, respectively. (**b**) We employed the built SG, PB, and PBSG models to predict RNA granule proteins with varying tiers from the RNAgranuleDB, respectively. (**c**) The average Gini feature importance of ten-fold validated models (N=10) was applied to select top ten most important features in the selected PBSG models (Gini feature importance results of SG and PB models are shown in **Fig. S9**). We performed our RNA granule models (*i.e.,* the selected PBSG model) to classify the collected SG proteomes (N = 253 in list 1^26^, N = 221 in list 2^7^, N = 486 in list 3^14^, and N = 172 in list 4^27^), excluded the training set proteins (**d**) or with overall proteins (**e**), respectively. Venn plots show the intersections between predicted positive proteins (*i.e.,* positive predictions) with collected RNA granule proteomes (**f**) and predicted non-RNA granule proteins (*i.e.,* negative predictions) in collected SG proteomes (**g**). HPI: hydrophobicity or hydrophilicity values. C: cysteine. W: tryptophan. L: leucine. LL: leucine-leucine. S: serine.

With the selected accurate and robust models, we proceeded to predict proteins with varying tiers from the RNAgranuleDB, as illustrated in **Fig. 2b**. We found the SG, PB and PBSG models could achieve similar prediction performance in terms of similar identified percentages (*i.e.,* 59.8% by SG models, 57.1% by PB models and 63% by PBSG models on PB tier 3 proteins in **Fig. 2b**) on SG, PB and PBSG tiers by our models, respectively. The consistent results among different RNA granule models on the RNAgranuleDB, also indicate the potential of our models to identify diverse RNA granule proteins. Notably, the percentages of different protein groups are consistent with the evidence standard and confidence (*e.g.,* from 100% of high-confidence tier 1 PBSG proteins identified to 56% of low-confidence tier 4 PBSG proteins identified). Moreover, the observed tendency aligns with existing six LLPS models (*i.e.,* PLAAC^29^, Cat Granule^30^, PScore^31^, DDX4-like^32^, R+Y^33^, and LARKS^34^) on identifying different tier proteins as LLPS-prone proteins, but with much higher identified percentages (*e.g.,* 42.9-100% by our models *versus* 16%^35^ by the six LLPS models) in the RNAgranuleDB. Therefore, compared with previous LLPS models with lower performance with lower PR AUC values and extensive identifications on the RNAgranuleDB, our RNA granule models could achieve much more reliable and more specific predictive performance on RNA granule proteins.

With the robust RNA granule models, our analysis gauged the roles of protein features within the constructed models in **Fig. 2c and S9-S11**. Cysteine residue fraction and gravy value are identified as the top two important features for RNA granule protein classification. Both show a negative relationship with RNA granule protein classification, indicating more fractions of cysteine and hydrophilic residues in proteins decrease the predicted propensity of proteins as RNA granule components in our models. Cysteine residues significantly influence protein 3D structures, primarily through disulfide bond formation, impacting protein folding^36^ and preventing from phase separation. Additionally, the extracted negative correlation from our models is also consistent with the notably lower fraction of cysteine residues in RNA granule candidates, compared to the human proteome and collected RBPs (**Fig. 1c, S2c and S3c**). The negative correlation emerges between gravy values with the RNA granule protein prediction probability within the human proteome highlights the significance of hydrophobicity in RNA granule protein classification. Consistently, related RNA granule literature reported that the presence of hydrophobic residues in IDRs of important RNA granule proteins (*e.g.,* FUS), as represented by the minus gravy values, plays a fundamental role in RNA granule formation, as previously suggested^12,37^.

Consequently, armed with these precise RNA models, we proceeded to scrutinize the connection between RNA granule protein probability (within the selected PBSG model in **Fig. S12**) and established domains or regions within selected highly assessed biomarkers for SGs (**Fig. S13 a&b**) and PBs (**Fig. S13c**). Our RNA granule models exhibit the capability to identify crucial domains (such as RNA recognition motif 2 (RRM2) in hnRNPA2B1, helicase ATP binding in DDX6, and Nudix Hydrolase in DCP2) and regions (comprising all IDRs in hnRNPA2B1, DDX6, and DCP2). These regions display relatively substantial prediction probability (exceeding 0.4) only within a sliding window of 30 residues. Our findings align with the established knowledge that certain domains, such as RRM, play pivotal roles in RNA granule formation^38^. Moreover, our results emphasize the importance of IDRs in proteins like DDX6 and DCP2, consistent with previous studies demonstrating their significance in LLPS and RNA granule formation^35,39^ in phase separation^39^. Furthermore, we observed that probability peaks tend to manifest alongside low gravy values and a low fraction of cysteine residues. This trend underscores the predominance of hydrophobicity and IDRs in RNA granule formation.

In addition, our objective extends to assess the model’s accuracy using four significant protein proteomes of SGs (N = 253 in list 1^26^, N = 221 in list 2^7^, N = 486 in list 3^14^, and N = 172 in list 4^27^), as depicted in **Fig. 2d-g**. Our RNA granule models effectively identified about 77% of the total proteins and over 60% of proteins, which were excluded in our RNA granule model training or testing. It is noteworthy that these SG proteomes were identified by different approaches (*e.g.,* proteomic analysis on the purified SG core^14^ and high-density proximity mapping^26^) under distinct experimental conditions (*e.g.,* yeast strain BY4741 at 30°C^14^ and human HEK293 cells at 37°C^26^) and diverse stresses (*e.g.,* heat shock stress^14^ and arsenite stress^26^). Meanwhile, only 14 in total 811 proteins (*i.e.,* less than 2%) were shared in all four published proteomes. Indeed, SG components were reported to be highly dependent on diverse types of stresses and cell types^7^, underscoring the challenges of our robust RNA granule models to have a comprehensive understanding on heterogeneous RNA granule proteome. As shown in **Fig. 2 f&g and S14**, the accurately identified SG components (77%) by our models belong to the component commonality among heterogeneous SG proteomes, while the unidentified SG components (23%) by our models come from the component diversity among heterogenous SG proteomes. It indicates that our RNA granule model holds promising potential to offer a comprehensive and holistic perspective for studying the critical RNA granule components contributing to the stability of their functionality independent of stresses.

Having established the comprehensive RNA granule proteome, we assessed the fundamental physicochemical patterns within the RNA granule proteome, as presented in **Fig. S15-S17**. The evident discrepancy in physicochemical patterns between the observed high-confidence RNA granule candidates sourced from the database with the identified RNA granule proteome, contrasted with the unlikely-LLPS PDB proteins and the identified non-RNA granule proteome from the human proteome, signals the compelling confirmation of formation patterns for RNA granules and substantiate the dependability of the robust machine learning models in grasping the multifaceted interactions that govern RNA granule classification. Simultaneously, this tendency in physicochemical patterns (*e.g.,* hydrophobicity) is also discernible within the identified RNA granule proteome across varying probability thresholds, as depicted in **Fig. S17**.

### Specific and Reliable Patterns in Identified RNA Granule Proteome to LLPS Proteome

In addition to the ongoing efforts to uncover the components of RNA granules, extensive research has focused on elucidating their functions, including posttranscriptional gene expression regulation^4^, mRNA translation^40^, RNA editing^9^, decay^9^, and storage^41^. In this study, we compared the enriched biological functions and domains of observed RNA granule proteins collected from the RNAgranuleDB with those predicted by our models and LLPS-prone proteins predicted by classic LLPS models^11,12,31^ in **Fig. S18, Fig. 3 and S19**. We could find a similar tendency between LLPS scores, RBP propensities and the predicted RNA granule probabilities derived from our RNA granule models in **Fig. S18**. Interestingly, RNA granule proteins exhibiting high likelihood with probabilities exceeding 0.9 corresponded to elevated prediction rank percentiles (over 0.8) across the three established LLPS propensity prediction models and high percentages as RBPs, as demonstrated in **Fig. S18 e&f and S20**. These findings suggest a strong correlation between the propensity for LLPS proteins, the propensity for RBP proteins and the likelihood of proteins to form RNA granules.

**Fig. 3.**
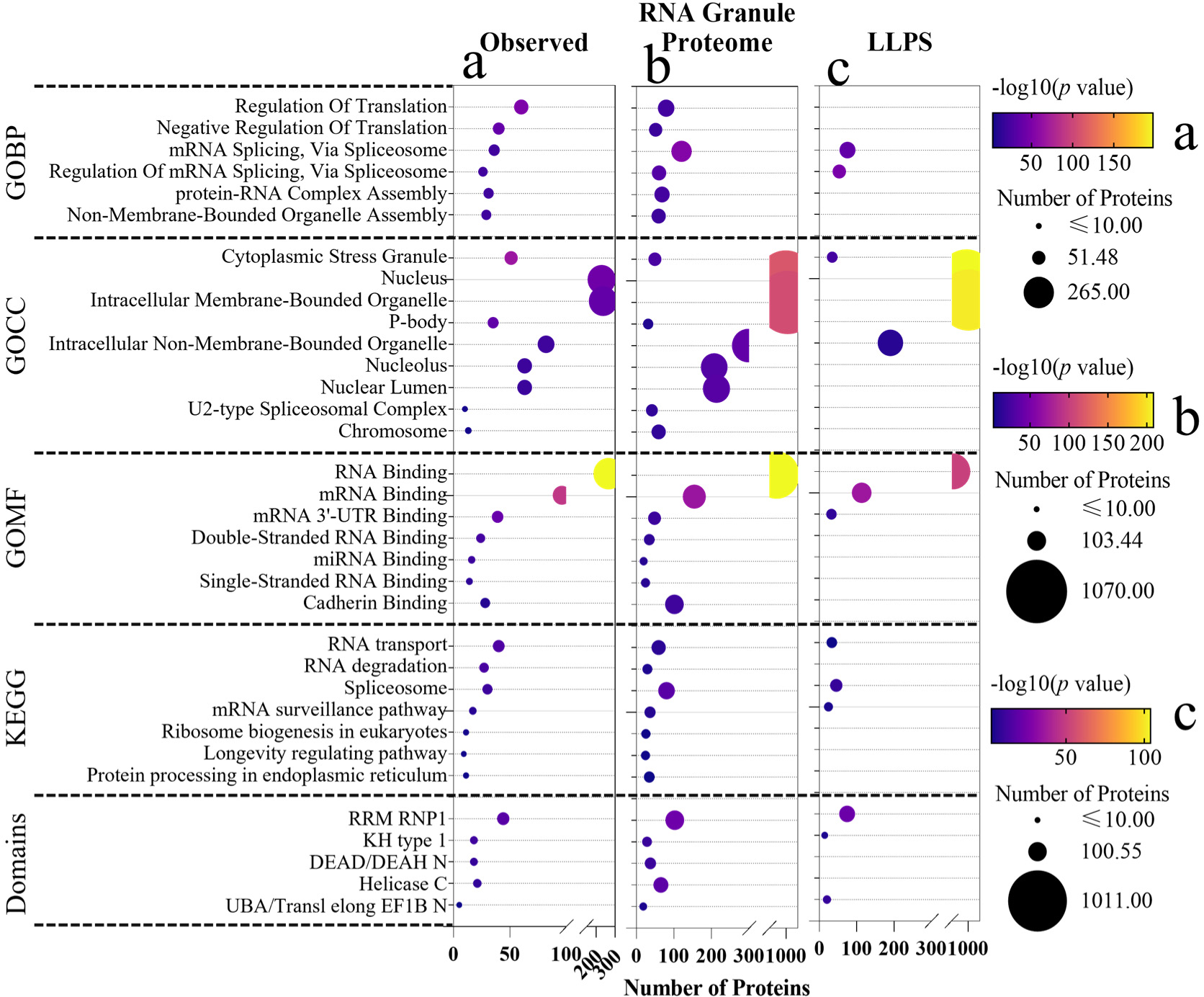
Functional enrichment analyses on the observed high-confidence RNA granule proteins (a, N=429), the high-confidence the identified RNA granule proteome (b, N=2225) and the predicted LLPS-prone proteome (c, N=2225). (**a**) We collected overall high-confidence RNA granule proteins utilized to train our models (*i.e.,* tier 1 for PBSG, N=429). (**b**) We selected the high-confidence RNA granule protein candidates according to the average prediction probability (over 0.7, the least of prediction probability of high-confidence RNA granule proteins by the selected PBSG models, N=2225) of the selected ten-fold evaluated PBSG models. Then, we completed the functional enrichment analysis on Gene Ontology biological process (GOBP), GO cellular component (GOCC), GO molecular function (GOMF), Kyoto Encyclopedia of Genes and Genomes (KEGG) and domains of the group of proteins. (**c**) We utilized three classic and widely applied LLPS prediction models (PSAR, DeePhase and PScore) to select high-confidence LLPS-prone proteins (N=2225) according to the average rank percentile values of prediction LLPS scores. The enrichment analysis is performed to compare the functional enrichment results of LLPS-prone proteins with identified and observed high-confidence RNA granule proteins. We applied the Enrichr platform to achieve the enrichment analysis.

As anticipated, the enrichment analysis presented in **Fig. 3&S19** reaffirmed the consistent pronounced enrichment of highly confident RNA granule candidates sourced from the database with enrichment of high-confidence and overall RNA granule proteomes. The consistent significant enrichment analysis indicates that RNA granule proteome identified by our RNA granule models, could show potentially related functions involved in similar biological processes, such as RNA transport^40^, RNA editing^9^, RNA decay^9^ and non-membrane organelle assembly (*i.e.,* a key process for the membraneless RNA granule assembly). Remarkably, our predictions revealed significant enrichment of these pivotal domains (*e.g.,* RRM RNP1, DEAD/DEAH N, and Helicase C) within the predicted RNA granule proteome from the human proteome. RRM RNP1 domains are known to be involved in RNA binding, which is a fundamental aspect of RNA granule function^42^. The observation further demonstrates the reliability of the identified RNA granule proteome by our machine learning models in terms of intricate biomolecular grammars in RNA granule assembly and functions.

In addition, the distinct differences in the top 20 significantly enriched functional terms and domains between predicted LLPS-prone proteins with observed high-confidence RNA granule proteins in **Fig. 3 c&a and Fig. S19 c&a**, respectively, highlight the limited capability of general LLPS models in the specific identification of RNA granule proteins.

### Evaluation of Community Grammars of Identified RNA Granule Proteome

To explore the community grammars within the predicted comprehensive RNA granule proteome, we constructed the RNA granule proteome community using physical PPIs grounded in experimental evidence from the STRING database^43^, as shown in **Fig. 4a**. Our analysis identified 6600 RNA granule proteins shown in the PPI network, averaging 46 interactions per protein. The human proteome PPI network comprises 19126 proteins and 947158 experimentally verified edges, averaging 50 PPIs per protein. To discern key nodes contributing significantly to this network, we employed seven metrics: degree, betweenness centrality^44,45^, eigenvector centrality^46^, PageRank^47^, closeness centrality^48^, clustering coefficient and degree centrality, as illustrated in **Fig. 4b-h**. It demonstrates a conspicuous increasing trend in all seven community importance metrics, ranging from low-probability to high-probability (*i.e.,* prediction probability from 0.5-1.0) RNA granule proteins. The observation aligns with previous experimentally identified RNA granule cores in a dense PPI network^7^, identified in specific experimental environments^7^. Notably, the high-confidence already known RNA granule proteins candidates comprise up to only 33% of proteins from prediction probability from 0.5-0.6 to 0.8-0.9, as listed in **Table S2**. The highly evaluated and popular SG proteins in diverse experimental SG proteomes also perform high centralities with dense interactions in our RNA granule community. For example, the YBX1^49,50^, AGO1, AGO2^51^ and UPF1^52^ present a central role in the RNA granule PPI network regard with 0.98, 0.95, 0.98 and 0.97 percentile rank of their PageRank values, listed in **Supplementary Files**. It indicates the centralities of these proteins beyond over 95% of proteins in the RNA granule PPI community (N=6600).

**Fig. 4.**
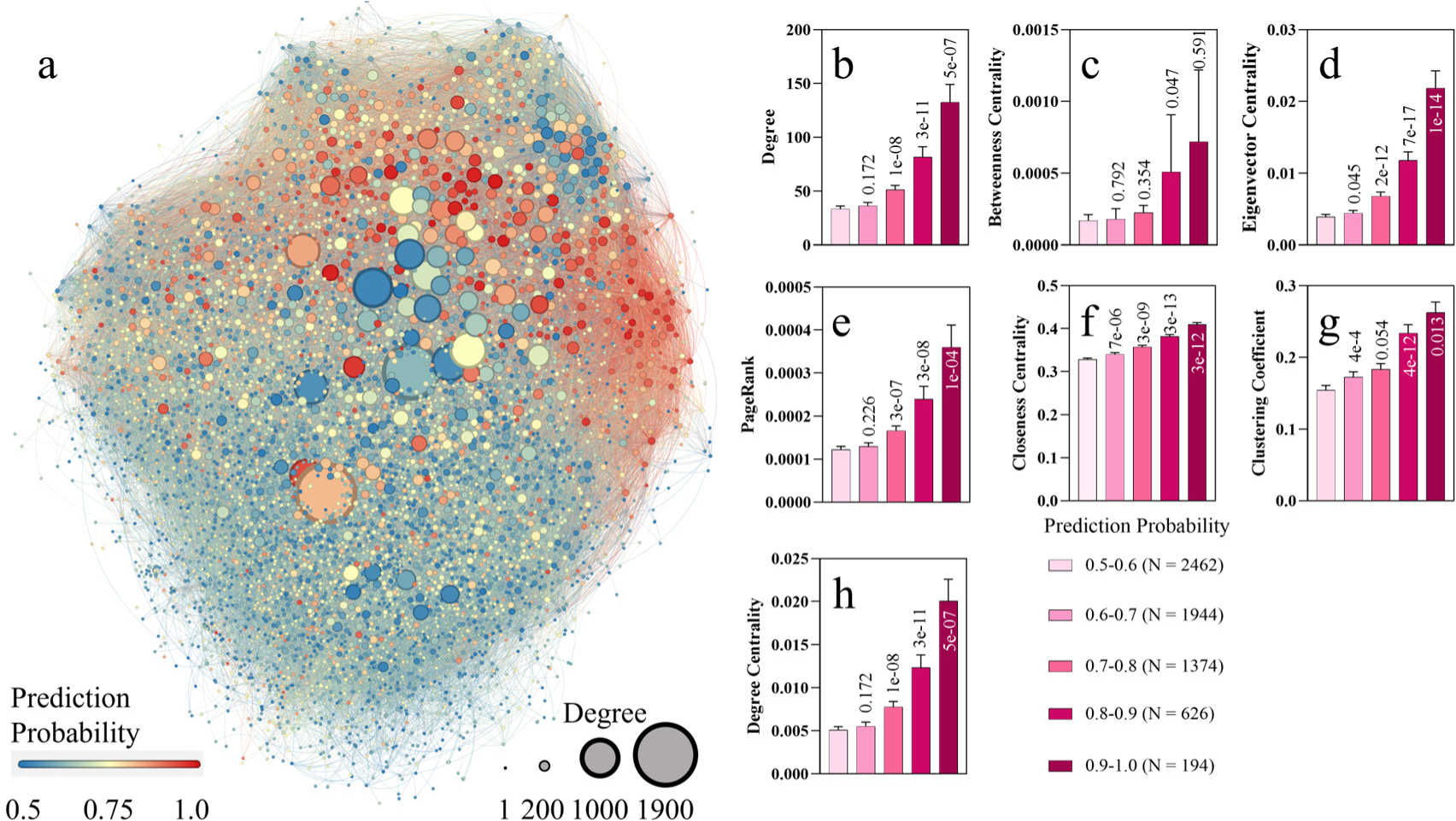
The centrality of identified high-confidence RNA granule proteins in RNA granule community. (**a**) The network of identified RNA granule protein (with average prediction probabilities over 0.5 by the selected PBSG models, N=6600) interactions was analysed using graph theory. A graph representation of the network was generated with nodes representing proteins and edges representing interactions between proteins. (**b-h**) Various measures of node importance were computed to identify key proteins within the network. These measures included degree (**b**), betweenness centrality (**c**), eigenvector centrality (**d**), PageRank (**e**), closeness centrality (**f**), clustering coefficient (**g**), and degree centrality (**h**). The *p* value of the one-way ANOVA test between the data in the recent group with the previous group data (*e.g.,* the *p* value is 0.172 compared degrees of proteins with prediction probabilities from 0.5-0.6 with proteins with probabilities from 0.6-0.7), was shown on/below the bar.

To further assess the generalization of biomolecular and community grammars in the RNA granule community generated from our model, we present the patterns of diverse RNA granules (typically classified as RNA granules, *i.e.,* PB^53^, SG^53^, ribonucleoprotein granule^54^, Cajal body^55,56^, P granule^57,58^, Chromatoid body^59^, nuclear body^60^ and Midbody^61^) and non-RNA granules (typically not classified as RNA granules, lipid droplets^62^, PML bodies^63^ and postsynaptic densities^64^) in **Table S3** and **Fig. S21**.

Generally, classic RNA granules, such as PB, cytoplasmic SG, ribonucleoprotein granule, Cajal body, and P granule, show a prevalence of proteins with high prediction probabilities (*i.e.,* over 0.7) as RNA granule proteins in our model and high centralities (measured by PageRank values) in the RNA granule PPI network in **Fig. S21**. Diverse RNA granules and non-RNA granules exhibit distinct fractions of predicted RNA granule protein components collected from the GO database (**Table S3**). For instance, a substantial percentage of protein components in PB (74%), SG (90%), ribonucleoprotein granule (74%), Cajal body (68%), and Chromatoid body (85%) are identified as RNA granule proteins by our RNA granule model. Moreover, most of these classic RNA granules’ protein components are identified as RBPs (49%-92%), while limited RBP components are observed in non-RNA granules (30%-50%). These results align with previous studies emphasizing the significance of RBPs as essential components of RNA granules^4^ and their association with post-transcriptional regulation^8^.

The analysis also reveals that well-studied RNA granule proteins, such as RC3H1, RC3H2, VCP, and G3BP1, are prominent and central proteins in the RNA granule PPI network across diverse RNA granules (**Fig. S21**), play pivotal roles in mRNA translation, mRNA stability^65^, and stress responses^65,66^; disassembling SGs associated with degenerative diseases^67^; and SG assembly^68^, respectively. The similarity in biomolecular and community patterns observed across widely studied RNA granules, such as PB, SG, and ribonucleoprotein granule, suggests common biomolecular and PPI mechanisms in their general formation and functions.

Given the existing limitations in techniques for the comprehensive identification of RNA granule components *in vivo* or *in vitro*, a comprehensive understanding of the RNA granule proteome from available GO databases remains challenging. To address this limitation and further assess the generalization of biomolecular and community patterns within dynamic and complex RNA granules, we present the prediction probability and community centrality of each protein in published SG proteomes across different experimental conditions, as illustrated in **Fig. S22**.

Like the patterns observed in **Fig. S21** for classic RNA granules, the four SG proteome lists exhibit clearly tend to include key RNA granule proteins characterized by high centrality and probability in the RNA granule community. However, due to the inherent complexity of dynamic SG proteomes in real conditions, these lists encompass a broader range of off-centre (*i.e.,* low centrality values in the RNA granule PPI community) and not high-confidence (*i.e.,* prediction probabilities < 0.7 by our RNA granule model) RNA granule proteins compared with SG in **Fig. S21b**. **Fig. S23a** highlights the high diversity and cell-type dependence of SG proteomes, with only 14 proteins shared among all four SG proteomes. Despite this complexity, consistent with the results in **Fig. S21**, popular and well-studied RNA granule proteins, including G3BP1, RC3H1, and RC3H2, emerge among the four SG proteomes.

Considering the heterogeneity of components in dynamic and complex RNA granules, the observed tendencies in highly popular RNA granule proteins with high centrality in the RNA granule community and RNA granule protein propensities from our RNA granule model persist among SG proteomes, as shown in **Fig. S23b**. These results underscore the generalizability of biomolecular and community patterns generated by our RNA granule model across diverse RNA granules.

To explore the relationship between the biological implications of RNA granule proteins and their centrality in the RNA granule PPI network, we conducted a GO enrichment analysis on identified RNA granule proteins with different centralities, as presented in **Fig. S24**. Proteins with higher centralities, exemplified by those with node importance percentile ranks exceeding 0.9 compared to those below 0.1, show a stronger connection with fundamental RNA granule processes, such as RNA binding and mRNA binding *versus* metalloendopeptidase activity and metallopeptidase activity. The analysis reveals a clear association between the centrality of identified RNA granule proteins and their biological roles in RNA granule formation and functionality. Consequently, the observation suggests that central components, exhibiting high prediction propensity and dense PPIs, may contribute to the generalization in the formation and functionality of RNA granules, while off-centre proteins, with low prediction propensity and limited PPIs, likely contribute specificity in the formation and functionality of RNA granules by interacting with other molecules and responding to environmental cues, as shown in **Fig. S25**.

### Dense PPI Clusters Perform Key Functional Subunits of Diverse RNA Granules

To unveil potential key RNA granule communities characterized by dense PPIs, we employed t-distributed Stochastic Neighbor Embedding (t-SNE)^69^ to visualize the complex RNA granule proteome PPI communities in 2-dimensional maps, as depicted in **Fig. 5&S26**. Notably, the observed paucity of proteins in these clusters within the ER suggests that their functional roles are more prominent in the cytosol and nucleus, aligning with previous observations regarding the major locations of RNA granules in the cytosol and nucleus^70^. As listed in **Supplementary Files**, the protein components of Cluster 1 and Cluster 2 exhibited high-confidence prediction probabilities (average prediction probabilities of 0.85 and 0.75) and node centralities (average percentile ranks of PageRank of 0.81 and 0.77), respectively. These results highlight that the proteins within these clusters possess high propensities as RNA granule components by our model and play pivotal central roles, with node centralities surpassing over 77% of the overall identified RNA granule proteins in the community.

**Fig. 5.**
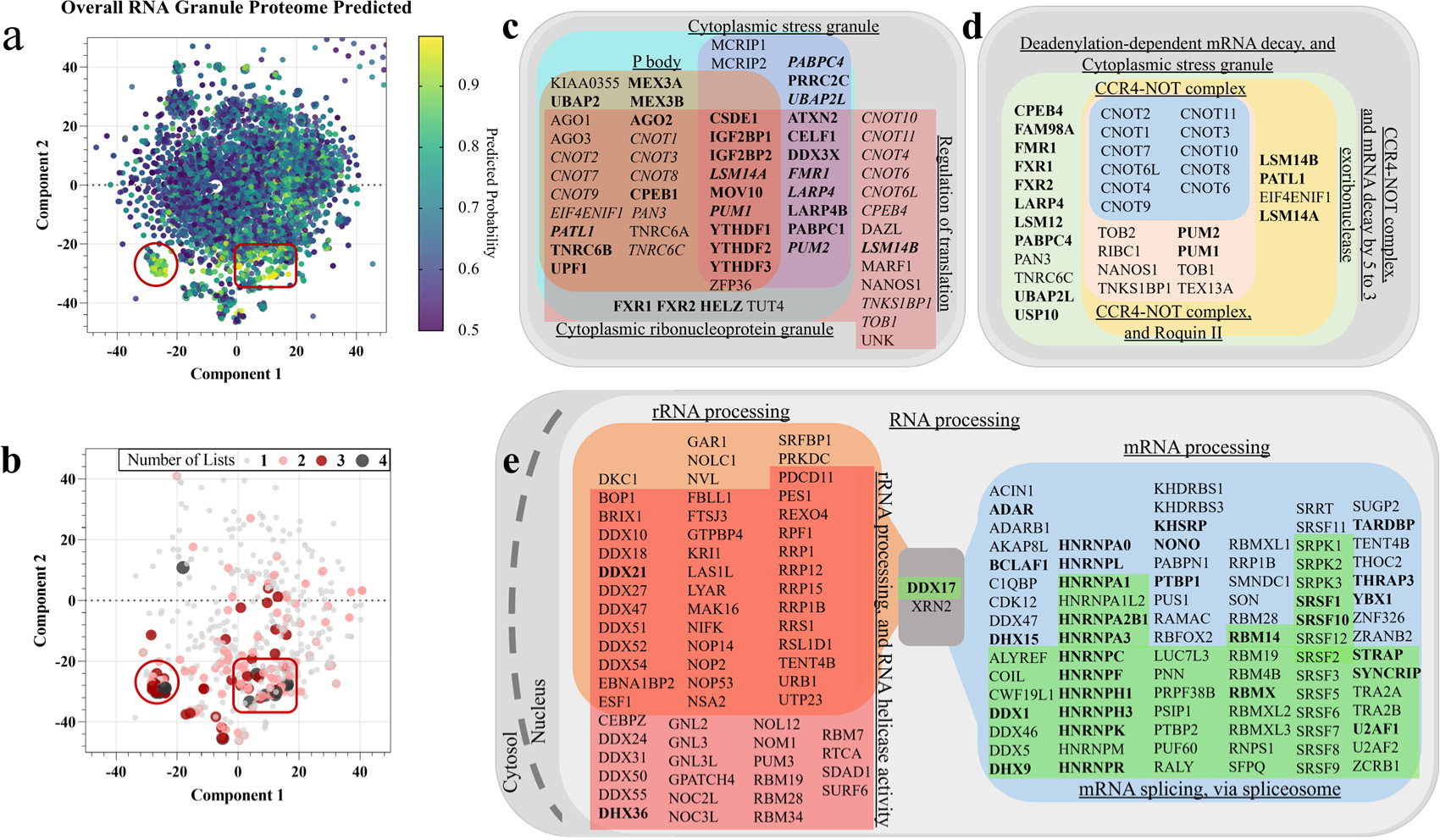
The generalization and function evaluation of extracted dense PPI clusters. (**a**) We visualized the overall identified RNA granule proteome PPI network (N=6600) into a 2-dimensional map and collected the location of each protein in the overall map (*i.e.,* the component 1 and component 2 values). (**b**) We visualized the common RNA granule proteins shared in different SG proteomes in the original RNA granule community. (**c&d**) We evaluated the potential biological implications by applying the GOCCs (*i.e.,* GO term: P body, GO ID: 0000932; GO term: Cytoplasmic stress granule, GO ID: 0010494; GO term: Cytoplasmic ribonucleoprotein granule, GO ID: 0036464), GOBP (*i.e.,* GO term: Regulation of translation, GO ID: 0006417 in **c**) and local network clusters (*i.e.,* term: CCR4-NOT complex, ID: 2931; term: CCR4-NOT complex, and Roquin II, ID: 2925; term: CCR4-NOT complex, and mRNA decay by 5 to 3 exoribonuclease, ID: 2924; term: Deadenylation-dependent mRNA decay, and Cytoplasmic stress granule, ID: 2922 in **d**) significantly (*p* value < 0.05) enriched by extracted proteins from Cluster 1. (**e**) We evaluated the potential biological functions by applying the GOBPs (*i.e.,* GO term: RNA processing, GO ID: 0006396; GO term: rRNA processing, GO ID: 0006364; GO term: mRNA processing, GO ID: 0006397; GO term: mRNA splicing, via spliceosome, GO ID: 0000398) and local network clusters (*i.e.,* term: rRNA processing, and RNA helicase activity, ID: 895) significantly (*p* value < 0.05) enriched by extracted proteins from Cluster 2. We applied the STRING platform to achieve the enrichment analysis. Bold: proteins occur in at least two of four SG proteomes from proteomic experiments. Italic: proteins also appear in the local network clusters of b. The circle represents Cluster 1. The square represents Cluster 2. ER, endoplasmic reticulum.

To assess the significance of the two identified clusters in RNA granules, we examined the distributions of these clusters within typically classified RNA granules and non-RNA granules (**Fig. S27&S28**). Our observations reveal the popularity of two clusters in diverse RNA granules, including PBs, cytoplasmic SGs, and ribonucleoprotein granules. In comparison with extensively studied PBs, SGs, and ribonucleoprotein granules, we observe that Cajal bodies, P granules, and Chromatoid bodies have been less comprehensively characterized, with smaller numbers of identified components from previous literature (N=60 for Cajal bodies, N=27 for P granules, and N=13 for Chromatoid bodies), listed in **Table S3**. Despite these limitations, even with a restricted number of identified protein components, these three typically classified RNA granules exhibit high percentages of components predicted as RNA granules by our model (up to 85%) and as RBPs (up to 92%), as demonstrated in **Fig. S21**. Unlikely, the protein components of these non-RNA granules tend to distribute outside the two selected clusters, particularly evident for lipid droplets and PML bodies. This observation aligns with the distinct formation and function mechanisms between lipid droplets^71^ and PML bodies^72^ compared to typically classified RNA granules, such as SGs.

Visualizing the protein distribution of the SG proteome identified through proteomic methods in **Fig. S29** reveals a notable localization of SG components from highly diverse experimental proteomes within the selected two dense clusters. This observation, coupled with the diverse components of the SG proteome depicted in **Fig. S23a**, where only 14 proteins (five of 14 proteins localized in Cluster 1 in **Fig. S30**; seven of them localized in Cluster 2 in **Supplementary Files**) are shared among all four SG proteome lists. **Fig. 5b** highlights that the two clusters encompass most of the common SG proteins. These findings strongly indicate that the two dense clusters may explain the stability and commonality of RNA granules across various stress and cellular contexts.

Exploring the enrichment analysis in the GOBP category in **Fig. S31**, Cluster 1 displays a significant association with biological processes closely related to RNA granule functions, such as negative regulation of translation and positive regulation of mRNA catabolic processes. These findings align with the known functions of RNA granules in translation, mRNA decay, and editing^9^. Moving to the GOCC enrichment analysis, proteins in Cluster 1 are enriched as components of diverse RNA granules, encompassing PBs, cytoplasmic SGs, and P granules. This observation further supports the notion that Cluster 1 is not only a component of various RNA granules but also plays a substantial role in their formation and function.

**figure. 5 c&d** reveals that many RNA granule proteins in Cluster 1 interact with the CCR4-NOT deadenylase complex^73^. The CCR4-NOT complex is observed in RNA granules, such as well-studied PBs and SGs, containing mRNA decay machinery and translation initiation factors, respectively^74^. Meanwhile, there are several related local network clusters enriched by Cluster 1 components in the STRING platform, such as the CCR4-NOT complex; CCR4-NOT complex, and Roquin II; CCR4-NOT complex, and mRNA decay by 5 to 3 exoribonuclease. The several related local network clusters underscore the significance of dense PPIs among Cluster 1 components in the post-transcriptional regulation of gene expression (*e.g.,* mRNA decay).

Unlikely, there are two connected dense subcommunities emerged within popular Cluster 2 (**Fig. S32**), housing numerous popular RNA granule proteins pivotal for RNA granule formation and functions, including notable protein family such as the DEAD box protein family (*e.g.,* DDX51, DDX27, DDX10, DDX18, and DDX31) aligning with one subcommunity, while the HNRNP family (*e.g.,* HNRNPA3, HNRNPF, HNRNPA0, HNRNPC, and HNRNPK) populating another. Notably, there are most of protein components (*i.e.,* 262 in 351) in Cluster 2 predicted by our models rather than already known RNA granule components in the training set, listed in **Table S4**.

Specifically, DEAD box proteins promote rRNA processing and transcription^75–77^. This observation is consistent with GO enrichment analysis results highlighting the significant enrichment of RNA binding and rRNA processing GOMF and GOBP terms, respectively, in **Fig. S33**. Moreover, the hnRNP family members tend to appear in another subcommunity of Cluster 2 in **Fig. S32**. The hnRNPs are considered essential ribonucleoproteins, pivotal in RNA metabolism through mRNA splicing^78–81^, increasing mRNA stability, and regulating translation^82^. As shown in the Enrichment analysis of Cluster 2 (**Fig. S33**), we could also find the related GOBPs with rRNA splicing (*e.g.,* mRNA splicing, via spliceosome and regulation of mRNA splicing, via spliceosome) are significantly enriched by Cluster 2 components. Additionally, some hnRNP proteins are identified as members of the spliceosome complex, a complex machinery governing alternative pre-mRNA splicing^83^. Notably, proteins like DDX17 and XRN2 link the two subcommunities with dense PPI connections (**Fig. 5e&S32**). Multifunctional DDX17 and XRN2, participate in various biological processes, including rRNA processing^84–86^ and pre-mRNA splicing^87,88^. Their intricate roles provide a plausible explanation for its robust connections with two dense functional subcommunities within Cluster 2 of our RNA granule PPI network.

In alignment with our discussed findings, our exhaustive analysis of the RNA granule proteome community provides a novel perspective, elucidating connections between the identified RNA granules and their pivotal biological roles under realistic conditions. We present a scientific hypothesis regarding functional subunits in RNA granule formation and functions, as shown in **Fig. 6**. It proposes that dense PPI networks form distinct subunits with fundamental functions, centrally interacting with other RNA granule components in relatively stable cores, thereby contributing to the stability of specific functions within diverse RNA granules.

**Fig. 6.**
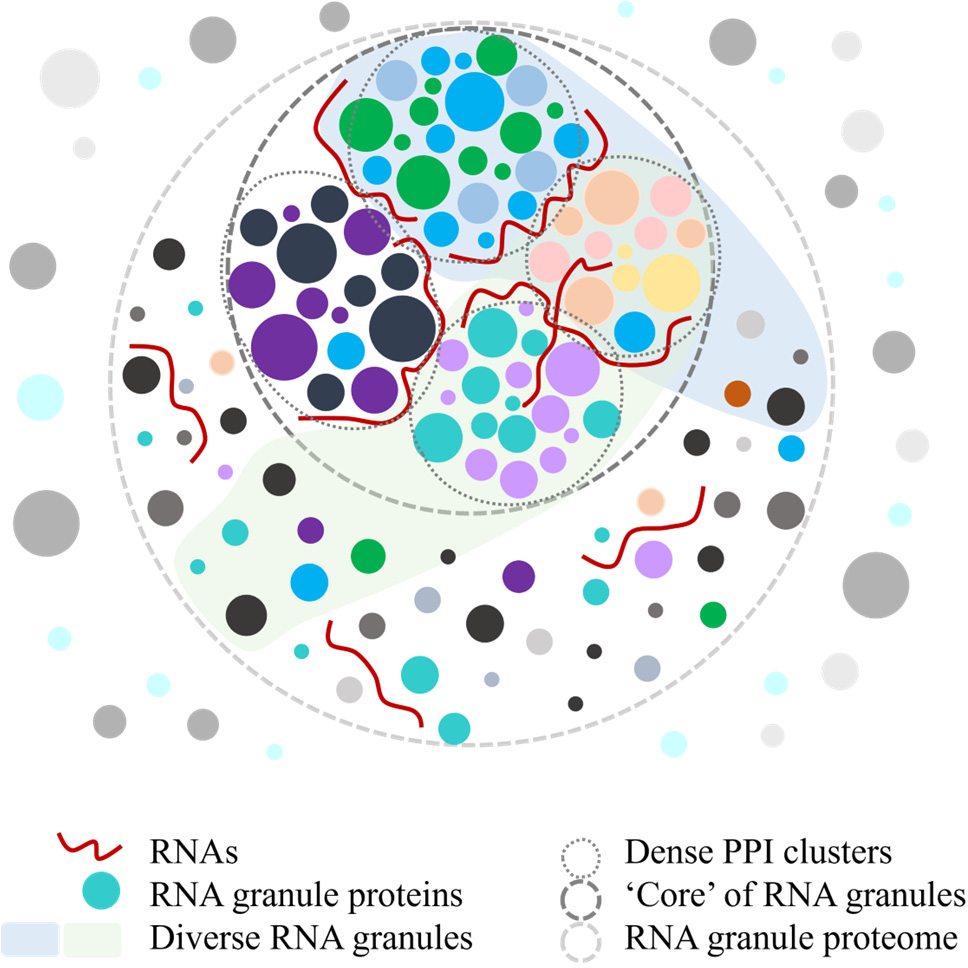
The hypothesis about RNA granule formation and functionality via functional PPI subunits. Our analysis reveals that dense PPI clusters, exemplified by Cluster 1 (associated with translation and mRNA decay) and Cluster 2 (linked to rRNA processing and mRNA splicing), form relatively stable ‘cores’ within diverse RNA granules, inspired by ^14^. These clusters significantly contribute to the stability of formation and functions of various RNA granules under distinct conditions.

## Discussion

This study employs machine learning methodologies to identify RNA granule proteome robustly, with more reliable and more specific prediction performance on RNA granule proteins than previous classic LLPS models^11,12,29–34^. Notably, the cysteine fraction and hydrophobicity emerge as the top two key features driving RNA granule formation, distinguishing the RNA granule proteome from both the human and the RBP proteome. Significantly, the predicted RNA granule proteome demonstrated substantial enrichment in biological functions associated with well-documented roles of RNA granules, such as RNA transport, spliceosome activities, and pivotal protein domains. This consistency aligns with the enrichment of functional terms and domains observed in the limited set of high-confidence RNA granule proteins documented within the database. These findings indicate the potential of our approach in constructing a comprehensive RNA granule proteome with high-confidence attributes.

The complex dynamics and heterogeneity of membraneless RNA granules, containing a myriad of RNAs and RBPs^8^, raise intriguing questions about their capability to maintain stability in post-transcriptional regulation and exhibit specificity in response to various stresses. Past experimental studies have unveiled the diverse components of RNA granules in response to diverse stresses. While traditional experimental methods identified key components of RNA granule proteomes under specific conditions^35^, comprehending the realistic mechanisms governing complex RNA granules requires an evaluation within a community context.

Our study employed the accurate RNA granule model to unveil PPI community grammars within the identified RNA granule proteome. Incorporating biomolecular grammars, such as prediction propensity from our RNA granule model, our analysis focuses on the commonality among typically classified RNA granules, non-RNA granules, and diverse SG proteomes across different conditions. Intriguingly, we note that typical RNA granules share similar biomolecular (*i.e.,* prediction propensities of proteins as RNA granule components by our RNA granule models) and community (*i.e.,* protein centralities in the predicted RNA granule proteome PPI community) grammatical patterns. Specifically, proteins predicted with higher propensities within RNA granules tend to occupy central roles in the overall PPI network RNA granules, underscoring the pivotal role of PPIs in the stability of formation and functions of RNA granules. This outcome concurrently validates the reliability of our RNA granule model.

Moreover, our analysis pinpoints two pivotal clusters marked by dense PPIs and high prediction propensities. Notably, these clusters exhibit significant sharing among diverse RNA granules and SG proteomes, transcending dependencies on stress or cell type. Through GO enrichment analyses, we uncover the involvement of these clusters in pivotal biological functions of RNA granules, including translation, mRNA decay, rRNA processing, and mRNA splicing.

In summary, our study sheds light on the pivotal role of PPIs in upholding the stability of the formation and functions of RNA granules. Offering a fresh and comprehensive perspective on the intricate nature of RNA granules, with a specific focus on PPIs, our findings emphasize the presence of shared functional subunits across diverse RNA granules. This analysis proposes a hypothesis: dense PPI clusters, exemplified by Cluster 1 for translation and mRNA decay, and Cluster 2 for rRNA processing and mRNA splicing, serve as integral functional subunits, constituting relatively stable ‘cores’ within diverse RNA granules. This contributes significantly to maintaining the delicate equilibrium between stability and specific functionality in the absence of membranes within the intricate cellular milieu.

## Methods

### Construction of the RNA Granule Dataset

As machine learning models are data-driven, the reliability and performance of the models largely depend on the dataset used. To construct our dataset, we retrieved samples of RNA granule proteins, including SG proteins, PB proteins, and PBSG granule proteins, from the RNAgranuleDB version 2.0 (http://rnagranuledb.lunenfeld.ca/, curation: November 2021 for SGs and February 2019 for PBs) as of February 2023^1^. Specifically, we focused on human proteins from the database, collecting all tiers (tier 1 to 3, or 4) for our SG, PB, or PBSG datasets, respectively. In addition, we collected all reviewed (Swiss-Prot) human proteins (20423 proteins) from the UniProt (https://www.uniprot.org/, Version: UniProtKB 2023_01) as of March 2023^89^. Subsequently, we filtered out proteins with sequence lengths less than 3 residues or truncated TTN protein to 32756 residues. Overall, we collected 20422 proteins from human proteome.

### Negative Data Construction

In the context of RNA granule research, previous studies have tended to report positive results, resulting in a shortage of negative samples in the RNA granule database and an imbalance between positive and negative samples for machine learning models. In this study, we considered all human proteins in the RNA granule database as positive candidates for our models. However, given that the RNA granule database may have missed some true RNA granule proteins among the remaining human proteins (excluding all RNA granule proteins in the RNAgranuleDB), we employed a random sampling strategy to construct negative samples with the same number of positive samples from the remaining human proteins to address this issue.

Based on previous LLPS models, the protein sequences in the publicly available PDB dataset^90^ have a low likelihood of undergoing phase separation and they were commonly used to be negative LLPS candidates for training relative models^12^. Therefore, the analysis also applied the PDB dataset from the work^12^ as negative samples to evaluate the performance of our specific RNA granule models.

### Physicochemical Features and Aa Content of Sequences

Inspired by the classic LLPS model^11^, we estimated important physicochemical properties and aa content of whole protein sequences and LCRs which have been proved to dominant LLPS. These physicochemical features included length, mw (Da), IEP, hydrophobicity (or hydrophilicity), aromaticity, Shannon entropy, cation fraction, protein secondary structures and low complexity. We applied the ProteinAnalysis module of Biopython package^91^ (Version: 1.80) in Python (Version: 3.8.8) to estimate the mw, IEP, gravy and aromaticity (*i.e.*, aromatic fraction) of protein sequences. The me values of all residues were summed up to estimate protein mw of each protein sequence. The IEP was calculated following the methods of Bjellqvist et al.^92,93^. The gravy values were calculated to evaluate the hydrophilicity of each sequence using the Kyte and Doolittle hydropathy scale^94^, which summed the individual hydrophilicity value of each residue in the sequence. In addition, we also estimated hydrophilicity by calculating total number or fractions of residues with hydrophilicity values lower than -1.5, -2 or -2.5 in each protein sequence, respectively. The aromatic fraction was calculated by determining the relative frequency of phenylalanine, tryptophan, and tyrosine in the whole sequence. The Shannon entropy of individual sequences was estimated using the formula^12^:

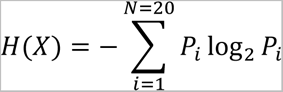

Where the variable *P* represents the frequency of the 20 naturally occurring residues in the given protein sequence. The cation fraction was estimated by calculating the relative frequency of lysine, arginine, and histidine in each protein sequence. As shown in the classic LLPS model, PSAP^11^, the fractions of protein secondary structures (*i.e.*, alpha-helix, beta-turn, beta-sheet) were estimated as the summed fractions of valine, isoleucine, tyrosine, phenylalanine, tryptophan and leucine (alpha-helix); asparagine, proline, glycine and serine (beta-turn) and glutamate, methionine, alanine and leucine (beta-sheet).

We defined and estimated the low complexity score by the number of unique residues in each sliding window (20 aas) as the PSAP^11^ model. The total low complexity scores or fractions were calculated as the number or fraction of aa within sliding windows (20 aas) with low complexity score ≤ 7 in each whole protein sequence, respectively. We estimated the low complexity score of each aa by counting occurrence of the amino acid in identified low complexity positions by sliding windows. The low complexity fraction of each aa was estimated by the ratio of the low complexity score of the aa to total number of collected low complexity windows. The lowest complexity score was estimated by the lowest complexity score of all sliding windows.

### K-mer Features for the Sequences

We extracted all available 2-mers and 3-mers from the human proteome and determined their fractions as the ratios of occurrence to each protein sequence length. To select important k-mers from human proteome, we counted the total occurrence of each k-mers in overall collected human RNA granule protein candidates from the RNA granule database and estimated the difference between k-mer fractions of collected RNA granule proteins with k-mer fractions of negative protein candidates randomly selected from human proteome with one-way ANOVA test using *Stats* module in *Scipy* package (Version 1.9.3) of Python (Version: 3.8.8). Then, we selected top significantly (*p*<0.001) 50 2-mers and 50 3-mers according to their occurrences among overall collected human RNA granule proteins from the RNAgranuleDB.

Overall, there are 179 features for each protein sequence in our model, including 60 features for fractions in each whole sequence, scores, and fractions in LCRs of 20 common residues, 3 features for the low complexity fraction, the low complexity score, lowest complexity score in the overall protein sequence, 16 features about physicochemical properties, 50 selected 2-mers and 50 selected 3-mers.

### Building of the Machine Learning Models

In this study, all RNA granule protein models were constructed using the *scikit-learn* package^95^ (Version 1.3.0) of Python. To achieve high accuracy and robustness, we employed the powerful ensemble machine learning algorithm, random forest^96^.

The random forest algorithm is a meta estimator that employs a collection of decision tree models on different subsamples of the dataset. By utilizing averaging techniques, a random forest can enhance predictive accuracy and avoid overfitting effectively^96^. We set the number of estimators as 2000, class weight as ‘*balanced*’ to employ class frequencies of the input data to automatically adjust weights and apply the ‘entropy’ criteria to evaluate the effectiveness of a split in a random forest model, inspired by the classic LLPS model^11^.

### Evaluation of the Machine Learning Models

To prevent overfitting of the models, we randomly divided the sampled dataset into ten folds to apply ten-fold cross validation to evaluate model’s performance. Meanwhile, we also applied the unlikely-LLPS PDB dataset as negative dataset to evaluate trained model’s performance. We also employed random seeds to ensure the reproducibility of our models, as demonstrated in the codes.

To evaluate the models’ predictive capability, we applied six important metrics in the analysis: accuracy, precision, recall, F1 value, the AUC and the PR AUC. The metrics were defined as below:

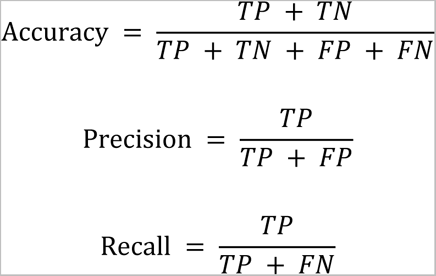

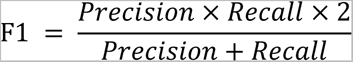

Where *TP* represents true positives, *TN* stands for true negatives, *FP* for false positives and *FN* for false negatives. Additionally, we utilized AUC values to assess the predictive performance of our RNA granule classification models. To determine the predictive performance at different possibility thresholds, we constructed ROC curves. By varying the threshold, we calculated the AUC values, which quantify the model’s performance on a scale from 0 to 1^97^. An AUC value of 1 represents a perfect prediction, while an AUC value close to 0.5 typically indicates a random guess. The PR AUC metric assesses the balance between precision and recall at varying decision thresholds. A higher PR AUC value signifies that the model is performing well in terms of achieving high precision while maintaining an acceptable level of recall. A PR AUC value of 1 represents a perfect prediction. The final RNA granule classification model applies the average prediction probability of ten trained models in the ten-fold evaluation to predict RNA granule protein scores of each protein. All metrics were calculated in the *scikit-learn* package (Version: 1.3.0) in Python (Version: 3.8.8).

### Sensitivity Analysis of Model Performance

The performance of individual protein models was influenced by various hyperparameters, including the number of protein features (*i.e.*, physicochemical features and k-mer features), the complexity of machine learning algorithms (*i.e.,* the number of estimators) and the number of sampled negative samples from the human proteome. To evaluate the sensitivity of model performance on the hyperparameters, we estimated models’ performance constructed with different protein features (*i.e.,* basic physicochemical features and aa contents, which were used in the classic LLPS model^11^, with 20 k-mers, 100 k-mers, 200 k-mers and all significant (*p* < 0.001) 837 k-mers), different numbers of estimators in random forest algorithm (*i.e.,* 1500, 2000 and 2500) and different ratios of sampled negative samples from the human proteome to positive samples (*i.e.,* 0.5, 1 and 1.5), respectively. .

### Feature Importance Evaluation

To evaluate the contribution of each feature in trained machine learning models, we estimated the Gini feature importance scores for each feature from ten trained random forest models in the ten-fold cross validation by using ‘*feature_importances_*’ function of ensemble module in *scikit-learn* package (Version: 1.3.0).

In the training processing of a random forest classification model, each node within the binary trees seeks the optimal split using the Gini impurity, which is a computationally efficient approximation to entropy^96^. This impurity measures how well a potential split separates the samples of the two classes at a particular node^96^. The Gini feature importance is essentially a by-product of this process, visualizing the outcome of the implicit feature selection performed by the random forest classification model and the significance of different features in the model^98^.

### Partial Dependence Evaluation

To visualize the relationship between key protein features with prediction propensity of an RNA granule protein candidate by our models, we applied the partial dependence method^99^ to estimate the average prediction propensity with actual feature values (*i.e.,* 5, 15, 25, 35, 45, 55, 65, 75, 85 and 95 percentiles) of each selected key feature in the whole human proteome. The method could show the way the selected feature affects the prediction propensity in the proteome level. The partial dependence method assumed the estimated key feature is independent and uncorrelated.

### Key Physicochemical Property Selection

The distance correlation, Pearson correlation and feature importance (as described in Feature Importance Evaluation) were applied to evaluate the strength, direction of the relationship and dependence between each physicochemical feature (*i.e.*, gravy, mw, hydrophobicity or hydrophilicity values (HPI) < -1.5, HPI < -2.0, HPI < -2.5, HPI < -1.5 fraction, HPI < -2.0 fraction, HPI < -2.5 fraction, alpha helix, beta turn, beta sheet, length, aromaticity, cation fraction, LCR fraction, IEP and entropy) with the target prediction probability of the selected PBSG model.

Distance correlation can detect both linear and nonlinear dependencies, and it is zero if and only if the variables are independent^100^. Pearson correlation, on the other hand, measures the linear relationship between two variables, and it varies from -1 to 1^101^. The distance correlation value was estimated by the python package *dcor* (Version 0.6), and the Pearson correlation was estimated by the stats module of the python package *SciPy* (Version 1.9.3).

### Sliding Window Analysis

We applied the sliding window method to evaluate our RNA granule model performance on key domains and IDRs determining phase separation capability and RNA binding propensity. There are three important biomarkers selected for the analysis, including hnRNPA2B1, DDX6 and DCP2. We set the sliding window as 30 residues and calculated the average prediction probability as RNA granule proteins of each sliding window by the ten-fold evaluated selected PBSG models. Then, we also calculated and showed the average values of the top two features (cysteine fraction and gravy) distributed on highly evaluated RNA granule biomarkers (*i.e.,* hnRNPA2B1, DDX6 and DCP2) within sliding window 30 residues.

### Model Performance on Published SG Proteomes

To evaluate the performance of our selected RNA granule models (for SG, PB and PBSG models), we collected published protein lists for SGs in papers (N = 253 in paper 1^26^, N = 221 in paper 2^7^, N = 486 in paper 3^14^, and N = 172 in paper 4^27^). The total number of collected published unique SG proteins is 811. We performed our RNA granule models (*i.e.,* the selected PBSG model) to classify the collected proteins (proteins excluded the model training set or overall proteins), respectively.

### Comparison with Classic LLPS Models

To evaluate the reliability of our identified RNA granule proteome, we applied classic and widely-utilized LLPS models to predict the LLPS propensity of human proteome, including the PSAP model^11^, the DeePhase model^12^ and the PScore model^31^. To evaluate the relative standing of each protein within all predicted LLPS proteomes, we calculated the percentile rank of each protein measuring the RNA granule propensities of each LLPS model and our RNA granule model (*i.e.,* the selected PBSG model). To visualize the relationship between RNA granule prediction propensities from our RNA granule model and LLPS propensities of all human proteome, we classified all human proteome into 6 groups in terms of RNA granule probabilities (*i.e.,* 0-0.5, 0.5-0.6, 0.6- 0.7, 0.7-0.8, 0.8-0.9 and 0.9-1.0) predicted by the selected PBSG model. Then, we visualized the distribution of LLPS propensities predicted by different LLPS models on each group of RNA granule proteins with different RNA granule probabilities, respectively.

Because the PScore model only predicts the probability of proteins with at least 140 residues. Therefore, we calculated the average percentile rank value of three LLPS models and selected top 6694 (the same number with predicted RNA granule proteome with RNA granule probability over 0.5) or 2225 (the same number with predicted RNA granule proteome with RNA granule probability over 0.7) proteins from human proteome (N=20422) according to the average percentile rank LLPS values. Then, we completed the enrichment analysis shown below and compared the functional enrichment analysis results of LLPS-prone proteins with overall RNA granule candidate proteins and collected high-confidence RNA granule proteins from the RNAgranuleDB.

### Enrichment Analysis in Enrichr

To assess the biological implications of the predicted RNA granule proteome and evaluate their reliability, we conducted KEGG pathway (2021 version), GO (2023 version) and the InterPro^102^ database (2019 version, access date July, 2023) enrichment analysis. This analysis aimed to identify enriched pathways in the KEGG database and GOBP, GOMF, GOCC and functionally important domains at the version. We utilized the observed high-confidence RNA granule proteins (*i.e.,* tier 1 proteins for SG, tier 1&2 proteins for PBs, and tier 1 proteins for combined PBSG), as well as the high-confidence or overall RNA granule proteome (with average probability over 0.7 or over 0.5) predicted by the selected RNA granule models, respectively. We compared the shared terms in top 20 KEGG pathways, GOBPs, GOMFs, GOCCs and domains enriched by observed high-confidence RNA granule proteins (*i.e.,* tier 1 for SG and PBSG, tier 1 and tier 2 for PB), with identified RNA granule proteins (i.e., RNA granule proteins with prediction probability >= 0.5 or >= 0.7) and selected LLPS-prone proteins, based on the -lg(*p* value) of the each term. The *p* value is calculated using the Fisher exact test, which is a statistical test for proportions that assumes a binomial distribution and independence of the probability of any gene belonging to any test, to assess the significance of observed differences between groups. For the enrichment analysis, we utilized Enrichr (https://maayanlab.cloud/Enrichr/), a web-based tool accessed on July, _2023_103–105.

### RBP Collection

The human RBPs (N=6163) were collected from the RBPbase database (https://apps.embl.de/rbpbase/, version 0.2.1 alpha, access date: Oct. 2023). We identified the human RBPs from the RBPbase database by combining 35 lists (Hs_HEK293-RIC, Hs_HuH7-RIC, Hs_HeLa-RIC, Hs_HeLa-RBDmap, Hs_K562-

serIC-chr, Hs_K562-serIC, Hs_HeLa-RNPxl, Hs_HEK293-pCLAP, Hs_microRNA- RIC, Hs_JURKAT-2018-RIC, Hs_JURKAT-2018-eRIC, Hs_HeLa-RICK-hc,

Hs_CARIC, Hs_Cyto-eRIC, Hs_Cyto-Ars-eRIC, Hs_Nuc-eRIC, Hs_Nuc-Ars-eRIC, Hs_OOPS_HEK293, Hs_OOPS_MCF10A, Hs_OOPS_U2OS, Hs_PTex_0015,

Hs_PTex_015, Hs_PTex_15, Hs_R-Deep, Hs_XRNAX-MCF7, Hs_XRNAX-HeLa, Hs_XRNAX-HEK293, Hs_XRNAX-MCF7-ars, Hs_HEK293-cRIC-SINV, Hs_HEK293-cRIC, knownRBPs-2014-Hs, SONAR-Hs, hasRBD-Pfam-Hs, RBPDB-

Hs, humanRBPs-2021).

### PPI Network Construction of Identified RNA Granule Proteome

To evaluate the role of identified RNA granule proteins in the protein community, we applied the human protein physical links of functional protein association networks in the STRING dataset (https://stringdb-downloads.org/download/protein.physical.links.full.v12.0.txt.gz, Version: 12.0, access date: July 2023). The STRING database integrates diverse sources of experimental evidence, co-expression data, and computational predictions to construct a comprehensive network of functional prediction associations^43^.

The analysis focuses on the human protein physical links with experimental proof of 947158 links and 19126 unique proteins. To extract the PPI community of identified RNA granule proteins, we focused on proteins with related links and applied prediction probability thresholds based on our RNA granule model. This model calculates the propensity of each protein to function as an RNA granule protein, with a prediction probability of 0 marking the lowest propensity and 1.0 indicating the highest. Specifically, proteins with a prediction probability greater than 0.5 were included in the overall RNA granule protein community (N=6600).

### Node Importance in PPI Network of RNA Granule Proteome

In this study, we employed graph theory to analyze the network of identified RNA granule protein interactions. A graph representation of the network using the Python (Version: 3.8.8) package *NetworkX* (Version 3.0)^106^, where nodes in the graph represented proteins and edges represented interactions between proteins.

To identify key proteins with pivotal roles in the BPs under investigation, various metrics of node importance were computed. The measures of node importance included the following: Degree: counts the number of interactions of each protein using ‘*degree*’ module of *NetworkX* package; Degree centrality: quantifies the number of interactions a specific protein has with other proteins using ‘*degree_centrality’* module of *NetworkX* package; Betweenness centrality: measures the extent to which a protein acts as a “bridge” between different parts of the network using ‘*betweenness_centrality’* module of *NetworkX* package^44,45^; Eigenvector centrality: measures the influence of a protein within the network, taking into account its direct and indirect connections using ‘*eigenvector_centrality’* module of *NetworkX* package^46^; PageRank: like eigenvector centrality, PageRank measures the influence of a protein within the network, taking into account its direct and indirect connections using *‘pagerank’* module of *NetworkX* package^47^; Closeness centrality: measures how close a protein is to all other proteins in the network using ‘*closeness_centrality’* module of *NetworkX* package^48^; Clustering coefficient: measures the extent to which a protein’s interaction partners also interact with each other in the unweighted network using ‘*clustering’* module of *NetworkX* package.

### Protein Component Collection of RNA Granules and Non-RNA Granules

In this analysis, we systematically collected protein components from typically classified RNA granules, employing defined terms and GO annotations for precise categorization. The selected RNA granules include: PB (term: P-body; GO: 0000932), SG (term: cytoplasmic stress granule; GO: 0010494), Ribonucleoprotein granule (term: cytoplasmic ribonucleoprotein granule; GO: 0036464), Cajal body (term: Cajal body; GO: 0015030), P granule (term: P granule; GO: 0043186), Chromatoid body (term: Chromatoid body; GO: 0033391), Nuclear body (term: nuclear body; GO: 0016604) and Midbody (term: Midbody; GO: 0030496).

Additionally, we collected protein components from three biocondensates not typically classified as RNA granules, serving as a comparison set. These biocondensates include Postsynaptic density (term: postsynaptic density; GO: 0014069), Lipid droplet (term: lipid droplet; GO: 0005811) and PML body (term: PML body; GO: 0016605).

The protein components were retrieved from the QuickGO database^107^ (https://www.ebi.ac.uk/QuickGO/, version 2023-10-06, access date: Oct. 2023). **Table S3** summarizes the collected components, facilitating a comprehensive comparison between RNA granules and non-typical RNA granules. This collection serves as a crucial foundation for subsequent analyses aimed at elucidating the distinct characteristics and functional roles of RNA granule protein components.

### PPI Network of RNA Granules in Different Cellular Locations

In the assessment of community grammars (*i.e.,* node importance of each protein) with the RNA granule proteome PPI network across various cellular locations, we compiled human protein from three crucial cellular compartments: cytosol (term: cytosol; GO: 0005829; N=2652), nucleus (term: nucleus; GO: 0005634; N=2752) and ER (term: endoplasmic reticulum; GO: 0005783; N=325). The comprehensive protein selection was obtained from the QuickGO database ^107^ (https://www.ebi.ac.uk/QuickGO/, version 2023-10-06, access date: Oct. 2023).

Then, we applied the extracted community properties (*i.e.,* PageRank value of each protein and percentile rank of PageRank value of each protein in the overall RNA granule proteome PPI network) of the collected proteins in three cellular locations in the whole RNA granule PPI network.

### t-SNE Visualization

In order to efficiently unravel the intricate structure of the RNA granule proteome PPI network, we employed the t-SNE technique^69^. The primary objective of applying t-SNE was to visualize the high-dimensional network in a two-dimensional map, assigning each protein a two-dimensional location (component 1, component 2) for intuitive interpretation.

t-SNE is particularly well-suited for revealing complex local structures, such as clustered subcommunities, by representing the similarities between data pieces in the original or embedded high-dimensional space by using Gaussian joint probabilities or Student’s t-distributions^69^. In our analysis, the t-SNE technique was completed, and the resulting two-dimensional locations of each protein within the entire RNA granule proteome PPI network (N=6600), as identified by our RNA granule model, were extracted.

The t-SNE visualization was implemented using the *‘TSNE’* module of the *scikit-learn* package (Version: 1.3.0) in Python (Version: 3.8.8). We configured the ‘*n_components’* as 2, while keeping other parameters as their default values.

### Distribution of Published SG Proteome on PPI Networks

To assess the performance of our selected RNA granule model, we gathered protein lists for SGs from four published papers (N = 253 in paper 1^26^, N = 221 in paper 2^7^, N = 486 in paper 3^14^, and N = 172 in paper 4^27^). The total number of collected published SG proteins is 811. Notably, we classified these proteins using our RNA granule model, which was trained separately.

To further evaluate the propensity of each collected SG protein component as an RNA granule protein, we utilized the prediction probability provided by our RNA granule protein model, as shown in **Supplementary Files**. Subsequently, we visualized the local structures of the collected published SG proteomes in a two-dimensional map using protein locations on the whole RNA granule proteome PPI network by t-SNE method.

### Extraction of RNA Granule Clusters

To identify critical functional subcommunities within the RNA granule proteome PPI network, we focused on two distinct groups of proteins exhibiting high prediction propensities as RNA granule proteins, as indicated by our RNA granule model. Subsequently, we extracted two clusters based on the two-dimensional locations of proteins within the entire RNA granule proteome community.

The criteria for cluster extraction were defined as follows:

Cluster 1: (-30.27 < component 1 < -23.70, -30.51 < component 2 < -21.42), N=100, as shown in **Supplementary Files** for detailed protein composition.

Cluster 2: (0.03 < component 1 < 19.80, -35.96 < component 2 < -20.00), N=351, as shown in **Supplementary Files** for detailed protein composition.

These criteria were established to capture distinct regions in the two-dimensional space potentially associated with functional subclusters within the RNA granule proteome.

### Functional Enrichment Analysis of RNA Granule Clusters

To evaluate the biological implications of extracted RNA granule clusters and assess their reliability, we performed functional enrichment analysis in Enrichr (https://maayanlab.cloud/Enrichr/), a web-based tool accessed on July 2023^103–105^ or in the STRING dataset (https://version-12-0.string-db.org/, Version: 12.0, access date: July 2023).

In the Enrichr, we conducted the GO (2023 version) enrichment analysis. We applied the analysis to identify enriched BPs, MFs and CCs with extracted clusters. We compared the top 10 significantly enriched BPs, MFs and CCs by extracted cluster proteins, based on the -lg(*p* values) of each term.

In the STRING, we constructed the PPI network for the extracted clusters with full STRING network. In the STRING networks, the edges between two proteins represent physical and functional protein associations and the network lines between two proteins indicate the strength of data support. We applied all available active interaction sources to construct the networks, including text mining, databases, experiments, co-expression, gene fusion, neighborhood and co-occurrence. Meanwhile, we establish a minimum required interaction score threshold of 0.4 (the default value).

The constructed PPI network for Cluster 1 comprised 100 nodes, 821 edges, an average node degree of 16.4, an average local clustering coefficient of 0.546, and a PPI enrichment *p* value < 1.0e-16. Similarly, the network for the extracted Cluster 2 included 351 nodes, 6163 edges, an average node degree of 35.1, an average local clustering coefficient of 0.511 and a PPI enrichment *p* value < 1.0e-16.

Subsequently, we conducted additional enrichment analysis on the PPI networks in the STRING database, encompassing GOBP, GOMF, GOCC and local network cluster (STRING) analyses.

## Statistics Analysis

All statistical analyses were completed using Python (version 3.8.6), GraphPad Prism (version 9.0.0) or OriginPro 2023b software (Learning Edition). The Pearson method was used to calculate all correlation coefficients unless specified otherwise, the one-way ANOVA in *SciPy* package^108^ was applied to calculated *p*-value, and all confidence levels of the interval were set to 0.95 unless specified otherwise.

## Data availability

The raw data and analysis code underlying this study will be made available upon requrest.

## Notes

### Competing Interest Statement

The authors have declared no competing interest.

