## Supplemental Information for "Unravelling biomolecular and community grammars of RNA granules via machine learning"

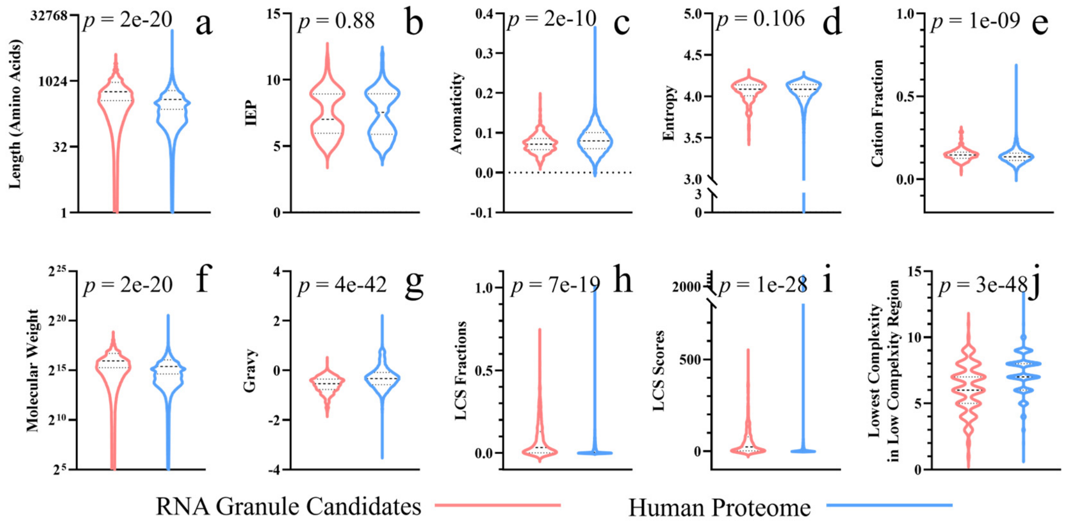


**Figure S1. Different RNA granule protein candidate and human proteome characterization on selected physical-chemical properties.** We compared the physico-chemical properties of observed high-confidence RNA granule protein candidates (*i.e.,* overall tier 1 proteins) from the RNAgranuleDB and the human proteome (excluded all RNA granule proteins in the RNAgranuleDB). We applied the one-way ANOVA to calculate the $p$ values.

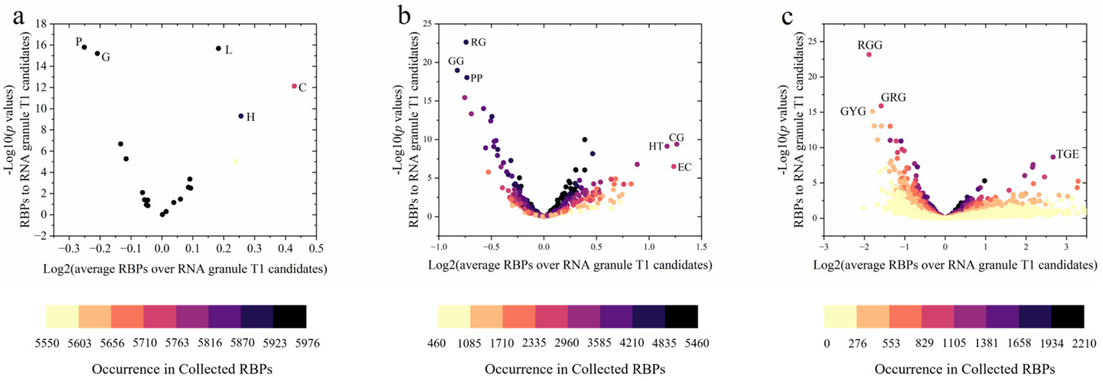


**Figure S2. Distinct k-mer fractions in RNA granule protein candidates (tier 1 proteins) compared to collected RBPs.** We compared the selected aa contents (a), 2-mer contents (b) and 3-mer contents (c) of observed RNA granule protein candidates and selected RBPs. To assess statistical significance, we applied one-way ANOVA to calculate *p* values comparing each k-mer fraction in RNA granule protein candidates with the corresponding fractions in collected RBPs. The occurrence counts the number of proteins containing the target k-mer among collected RBPs (N=6163). The fold change measures the ratios of the average fraction in collected RBPs to the average fraction in RNA granule protein candidates for each k-mer. P: proline. G: glycine. C: cysteine. L: leucine. H: histidine. RG: arginine-glycine. GG: glycine-glycine. PP: proline-proline. HT: histidine-threonine. CG: cysteinie. EC: glutamate-cysteine. GRG: glycine-arginine-glycine. RGG: arginine-glycine-glycine. GYG: glycine-tyrosine-glycine.

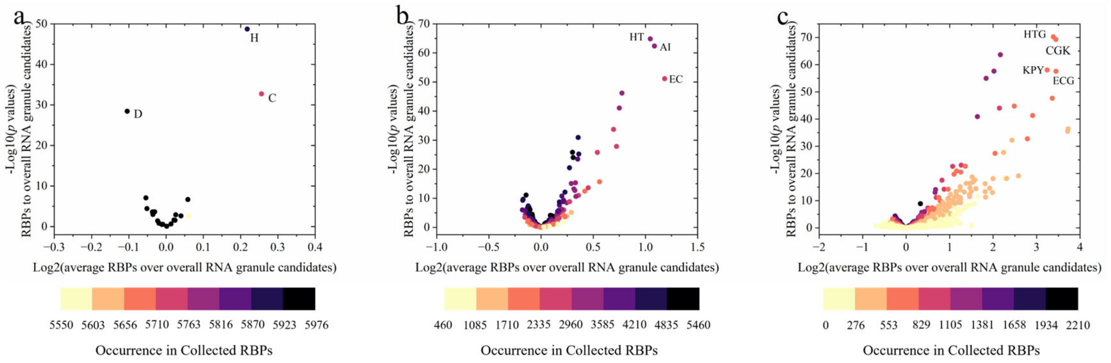


**Figure S3. Distinct k-mer fractions in RNA granule protein candidates (overall proteins) compared to collected RBPs.** We compared the selected aa contents (a), 2-mer contents (b) and 3-mer contents (c) of observed RNA granule protein candidates and selected RBPs. To assess statistical significance, we applied one-way ANOVA to calculate *p* values comparing each k-mer fraction in overall RNA granule protein candidates with the corresponding fractions in collected RBPs. The occurrence counts the number of proteins containing the target k-mer among collected RBPs (N=6163). The fold change measures the ratios of the average fraction in collected RBPs to the average fraction in overall RNA granule protein candidates for each k-mer. C: cysteine. H: histidine. D: aspartate. HT: histidine-threonine. AI: alanine-isoleucine. EC: glutamate-cysteine. HTG: histidine-threonine-glycine. KPY: lysine-proline-tyrosine. CGK: cysteine-glycine-lysine. ECG: glutamate-cysteine-glycine.

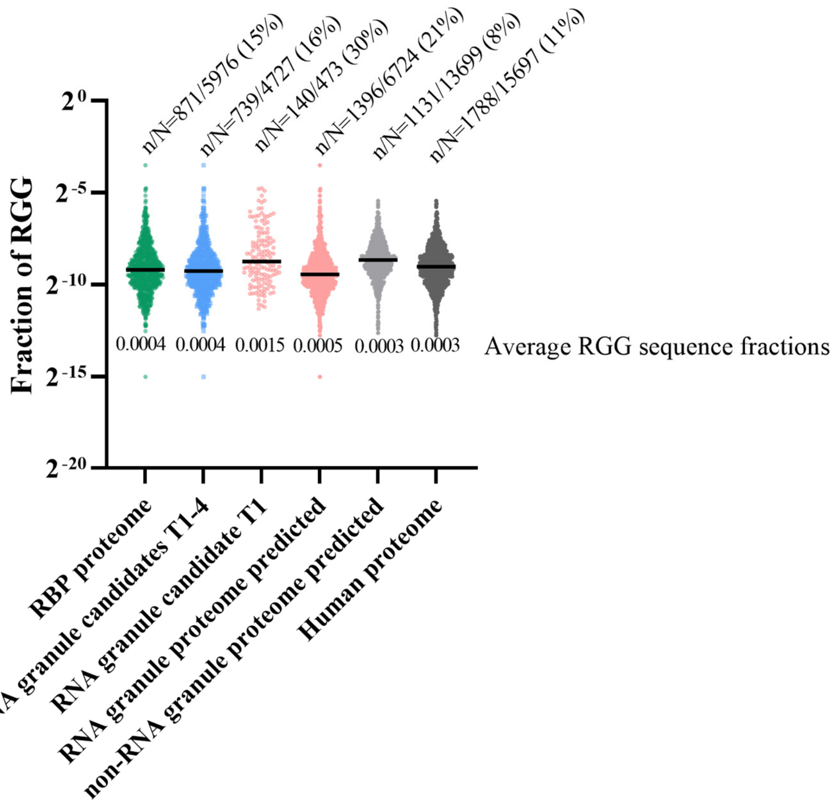


**Figure S4. Distinct RGG fractions in collected RBP proteome, overall RNA granule candidates (tier 1 to tier 4), high-confidence RNA granule candidates (tier 1), predicted RNA granule proteome, predicted non-RNA granule proteome and human proteome (excluded the overall RNA granule candidates).** We compared the contents of selected 3-mer, RGG, of proteins with RGGs in collected RBP proteome, overall RNA granule candidates (tier 1 to tier 4), high-confidence RNA granule candidates (tier 1), predicted RNA granule proteome, predicted non-RNA granule proteome and human proteome (excluded the overall RNA granule candidates). The average RGG sequence fractions measure the average fractions of RGGs in proteins of different groups. n: the number of proteins with RGG 3-mer in each group. N: the total number of proteins in each group.

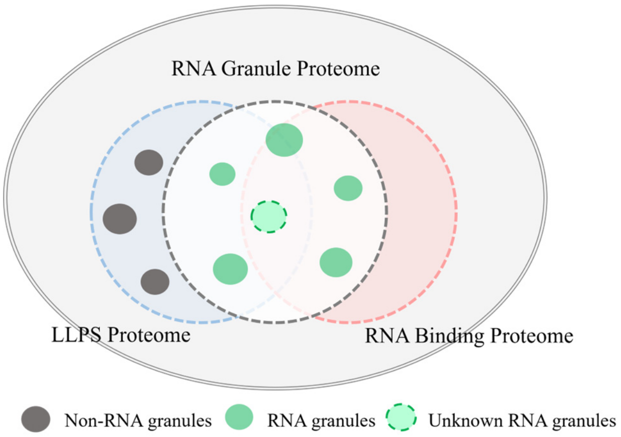


**Figure S5. The biomolecular mechanisms distinguish RNA granule proteome with LLPS proteome and RNA binding proteome.** As discussed above, the RNA granules tend to contain more protein components with large (*i.e.,* high mws) and hydrophobic residues (*i.e.,* protein gravy values < 0) compared with observed[1] and predicted[2] LLPS proteomes.

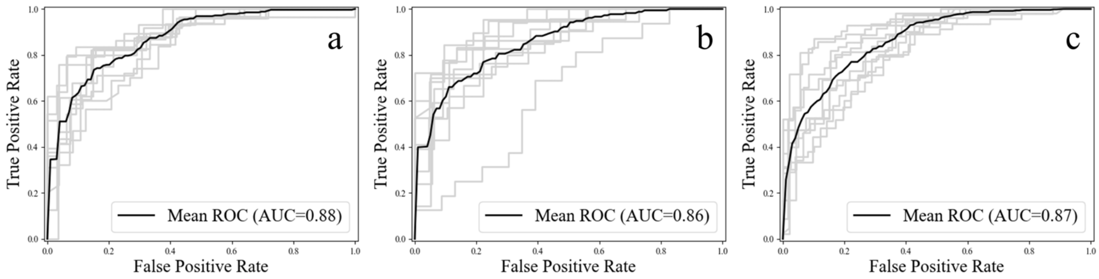


**Figure S6. Model performance of SG, PB and PBSG classifiers.** The analysis utilized ten-fold cross-evaluation to estimate the model prediction performance, quantified by the average AUC (represented by the black line) on testing sets for SG (a), PB (b), and PBSG (c) proteins. The gray lines depict the curves illustrating the ten-fold evaluation performance on each fold. True positive rate, also known as sensitivity, is defined as the likelihood of the model correctly predicting a positive outcome given that the actual outcome is indeed positive. True negative rate, also known as specificity, represents the probability that the model accurately identifies a negative outcome when the actual outcome is indeed negative.

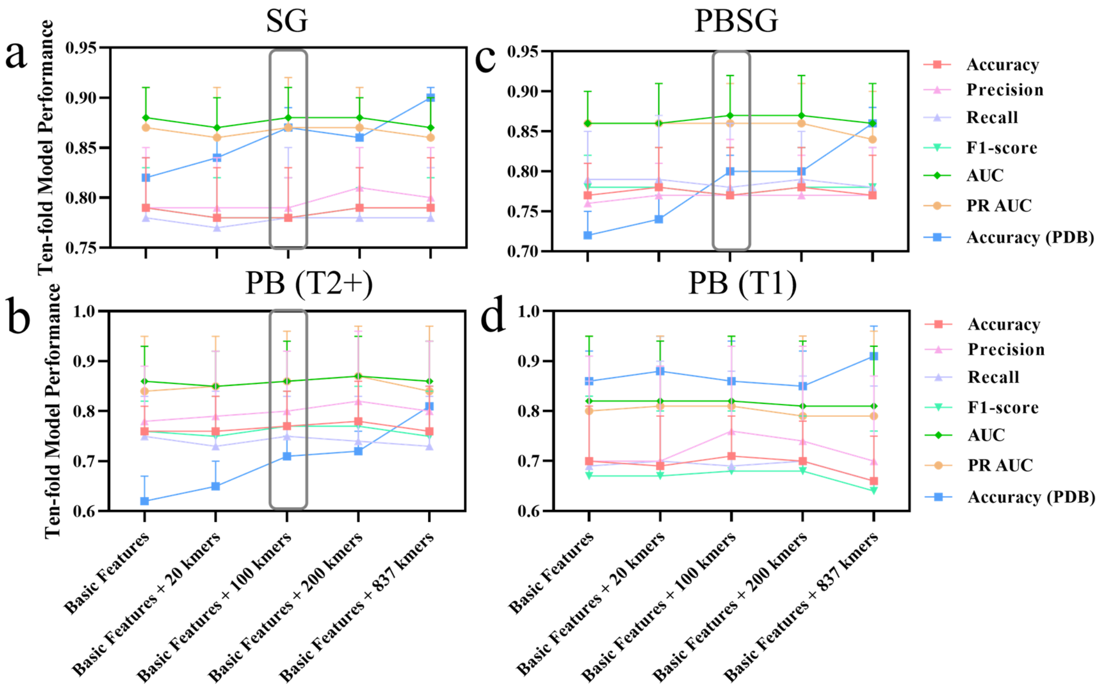


**Figure S7. Model performance evaluation on various 2-mer and 3-mer content features using high-confidence observed RNA granule proteins (tier 1 SG proteins in a, tier 1&2 PB proteins in b, tier 1 PBSG proteins in c, and tier 1 PB proteins in d) from the RNAgranuleDB, employing the ten-fold cross-evaluation method.** Basic features mean basic physico-chemical properties (N=19) and aa contents (N=40). The k-mers were selected according to significant difference ($p \leq 0.001$) compared to selected negative candidates from human proteome and abundance in observed RNA granule proteins. The terms '20,' '100,' '200,' and '837' k-mers refer to 2-mers (N = 10) and 3-mers (N = 10), 2-mers (N = 50) and 3-mers (N = 50), 2-mers (N = 100) and 3-mers (N = 100), and all 2-mers (N = 272) and 3-mers (N = 565) that show significant differences ($p \leq 0.001$) compared with negative candidates from the human proteome.

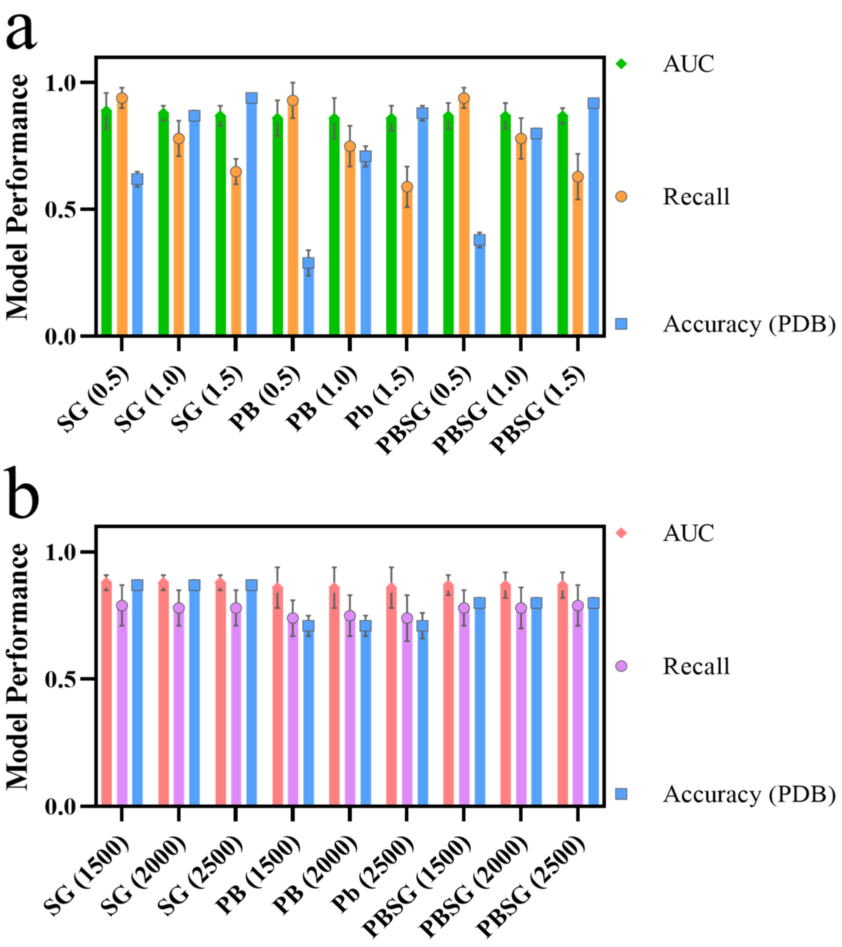


**Figure S8. Sensitivity analysis of model performance regarding the number of negative samples (a) and model complexity (b).** (a) In this analysis, RNA granule classifiers (SG, PB, and PBSG classifiers) were trained with different ratios (0.5, 1.0, and 1.5) of selected negative samples from the human proteome to the observed positive RNA granule proteins from the RNAgranuleDB. (b) In this analysis, RNA granule classifiers (SG, PB, and PBSG classifiers) were trained with different model complexities, varying the number of trees (1500, 2000, and 2500) in the random forest algorithm. Furthermore, we assessed the model performance by evaluating the AUC and recall on testing sets, as well as accuracy on collected unlikely-LLPS PDB proteins using ten-fold cross-evaluated models.

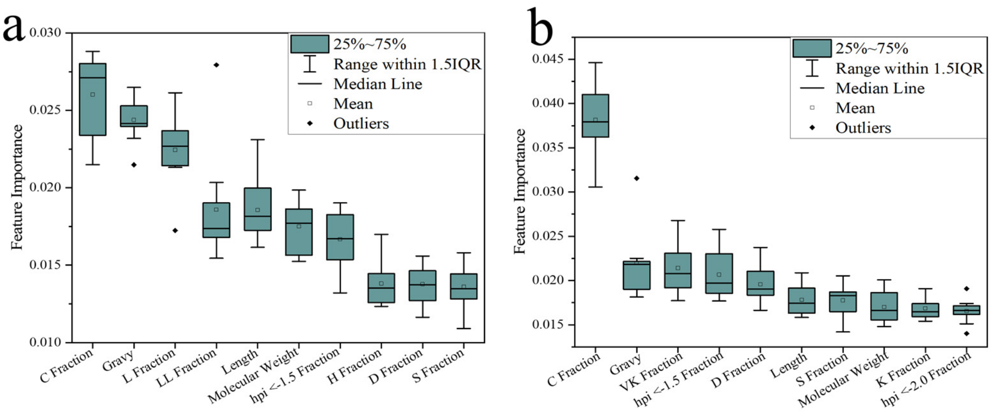


**Figure S9. Top ten important features of RNA granule classifiers (SG classifier in a and PB classifier in b).** The average feature importance of ten-fold evaluated models was applied to select the top ten most important features in the SG (a) and PB (b) classifiers, respectively. IQR: interquartile range.

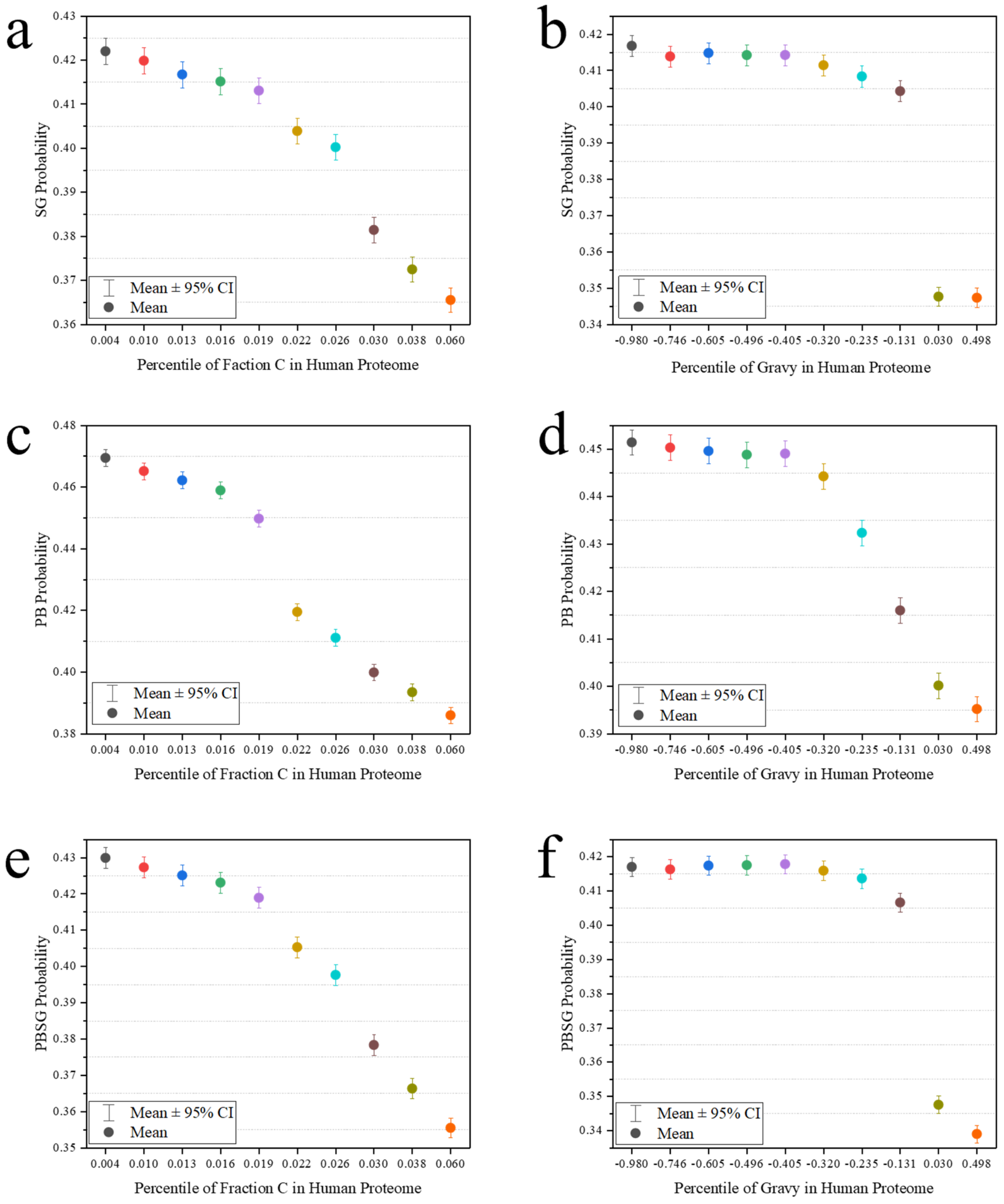


**Figure S10. Partial dependence shows the relationship between key features (fraction C and gravy) with prediction probability of SG, PB or PBSG classifiers, respectively.** The partial dependence method was employed to estimate the average prediction probability, considering actual feature values (*i.e.,* 5, 15, 25, 35, 45, 55, 65, 75, 85, and 95 percentiles) of the selected key feature in the whole human proteome data. The partial dependence method assumed the estimated key feature is independent and uncorrelated.

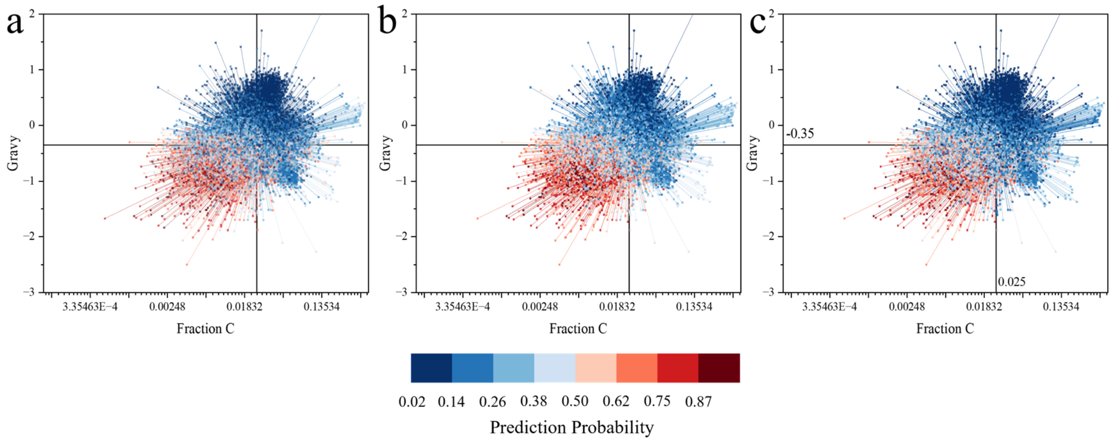


**Figure S11. Prediction probability of human proteome (SG in a, PB in b and PBSG in c) distribution on two key features (fraction C and gravy).** The centroid was performed by the Origin Pro software.

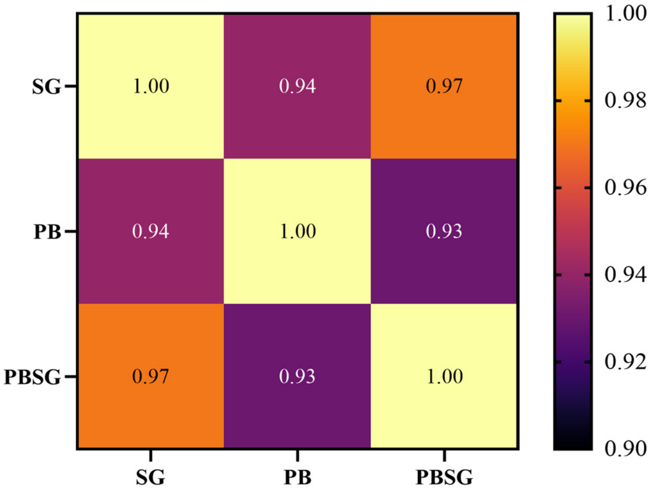


**Figure S12. The similarity of prediction probability of human proteome (N=20423) of three RNA granule classifiers (*i.e.*, SG, PB and PBSG classifiers).** The similarity was estimated by the Pearson correlation coefficient.

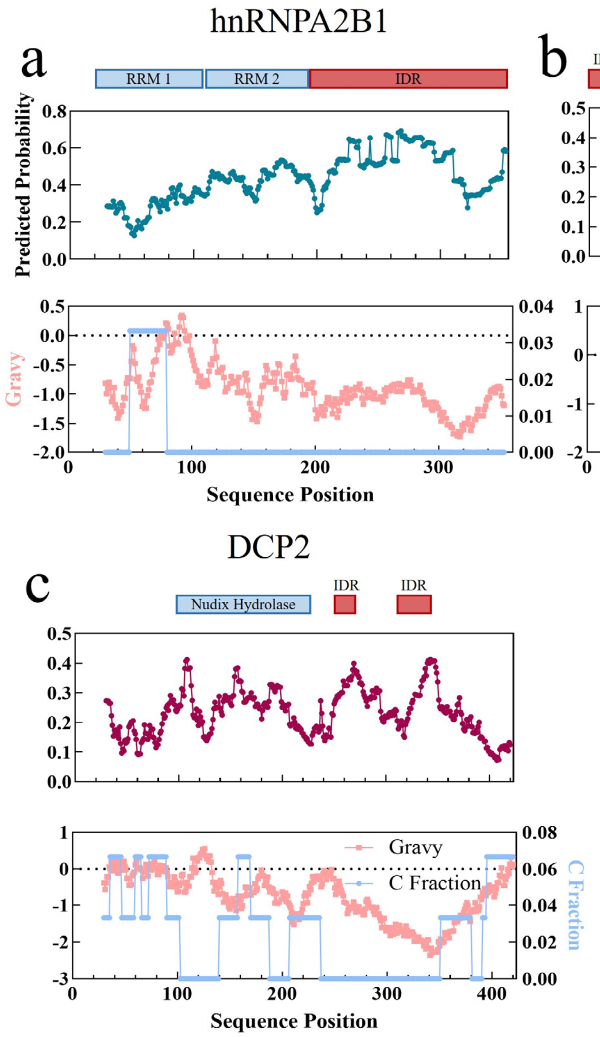


**Figure S13. model performance on highly evaluated RNA granule biomarker proteins.** The average prediction probability of the ten-fold evaluated selected PBSG classifiers was performed on highly evaluated RNA granule biomarkers (hnRNPA2B1 in a and DDX6 in b for SG; DCP2 in c for PB) with sliding window 30 residues. The average top two features (C fraction and gravy) distribute on highly evaluated RNA granule biomarkers (hnRNPA2B1 in a and DDX6 in b for SG; DCP2 in c for PB) with sliding window 30 residues.

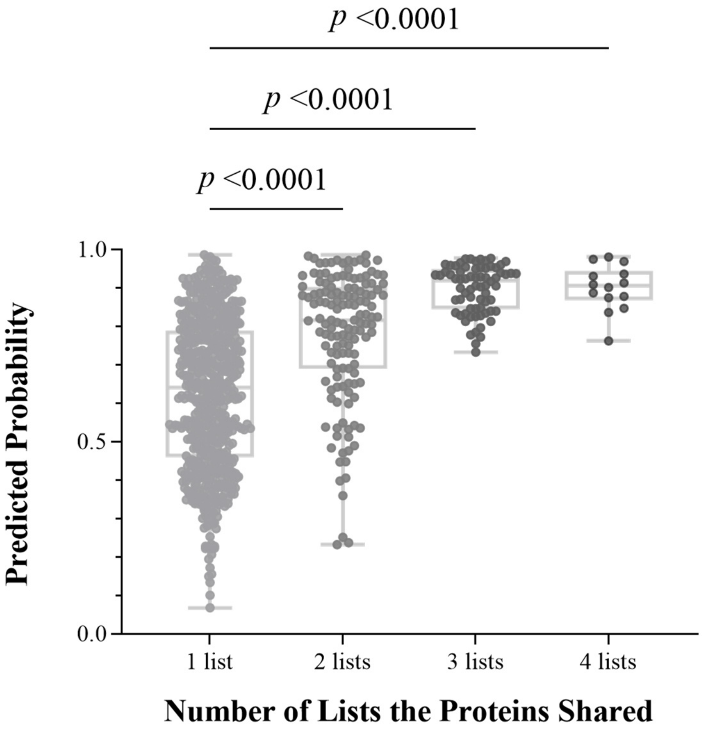


**Figure S14. Comparison on prediction performance of the RNA granule model on general and specific SG proteins among four published lists.** The one-way ANOVA analysis was applied to compare the prediction probabilities of proteins shared in only one list, two lists, three lists or four lists by the RNA granule model**.**

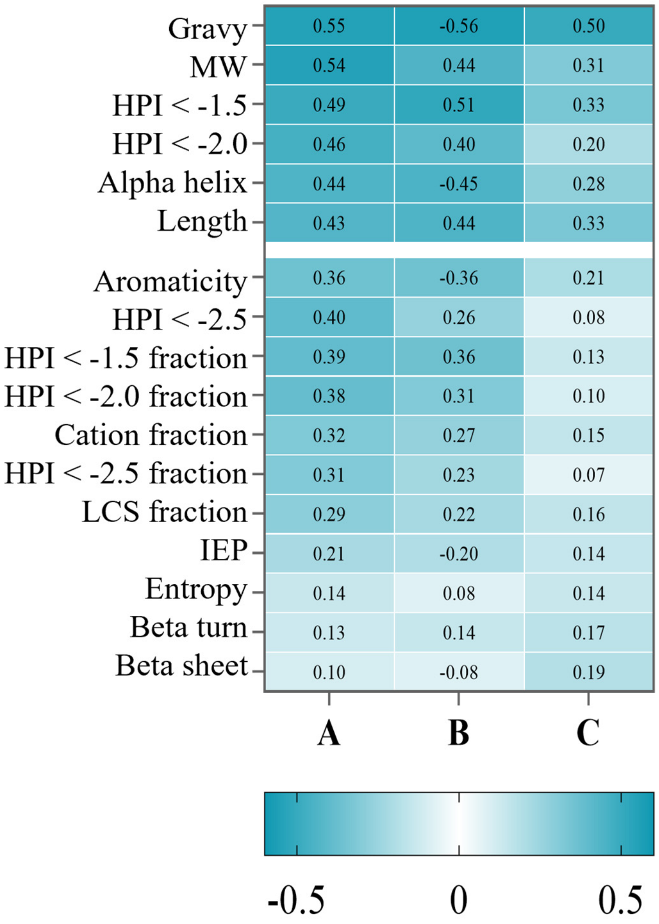


**Figure S15. The top six important features selected by distance correlation coefficient (A), Pearson correlation coefficient (B) between predicted propensities with selected protein features of the human proteome by our RNA granule classifiers, and mean feature importance*20 (C) in RNA granule classifiers.**

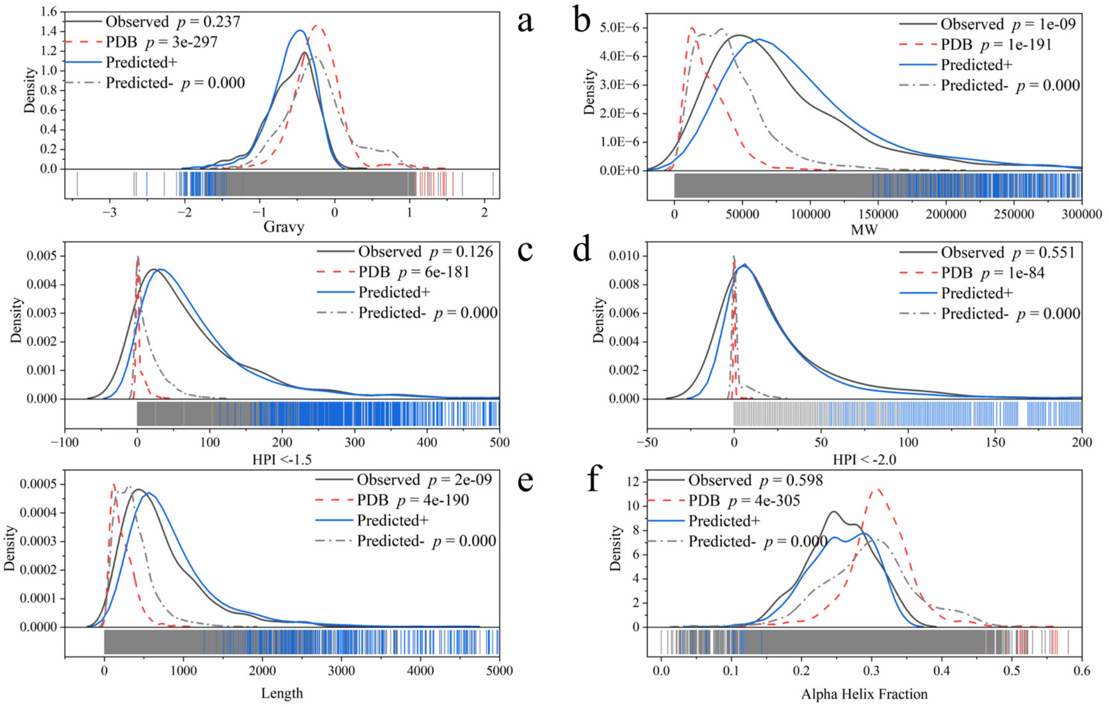


**Figure S16. Observed and predicted RNA granule proteins on selected important physico-chemical features.** We identified the RNA granule proteome from the human proteome with the selected PBSG classifiers with probability over 0.5. Observed data sets were high-confidence tier 1 or tier 2 proteins (tier 1 for SG, tier 1 for PBSG and tier 1 & 2 for PB) from the RNAgranuleDB, utilized to train and test RNA granule classifiers. We calculated the *p* value of the one-way ANOVA test between the predicted RNA granule proteome (with probability over 0.5) with the other group data (e.g., the *p* value is 0.237 compared the gravy values of predicted RNA granule proteome with observed high-confidence RNA granule proteins (tier 1 for SG and PBSG, tier 1&2 for PB) from the database).

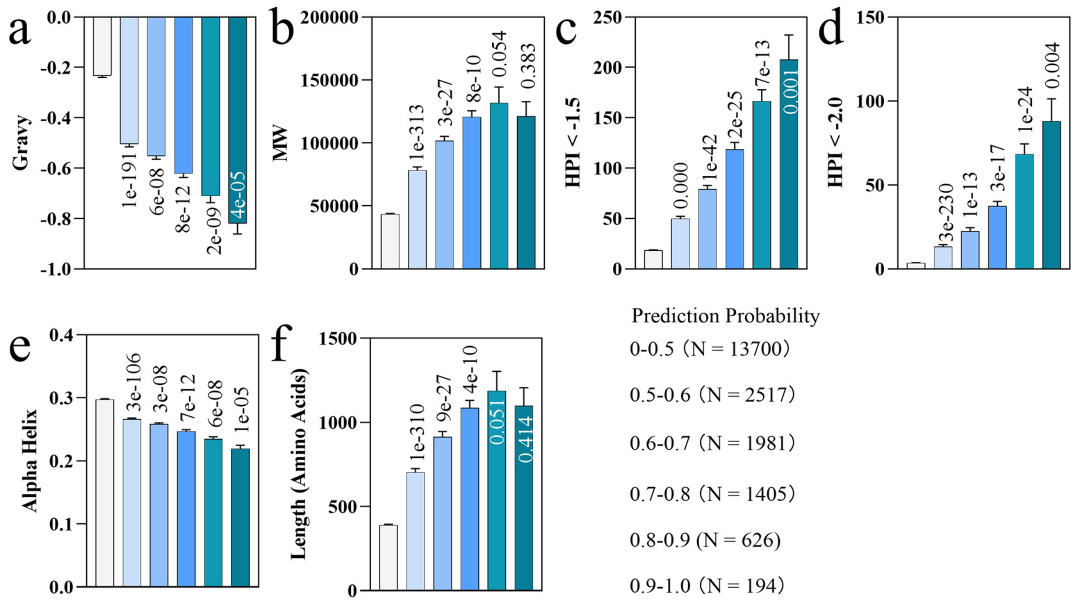


**Figure S17. Selected important physico-chemical features in identified RNA granule proteome from human proteome with different prediction probabilities by the selected PBSG classifier.** The bar represents the average value of each feature and error bars represent the 95% CI value of each feature. The *p* value of the one-way ANOVA test between the data in the recent group with the previous group data (*e.g.*, the *p* value is 1e-191 compared gravy of proteins with prediction probability from 0-0.5 with proteins with probability from 0.5-0.6), was shown on/below the bar.

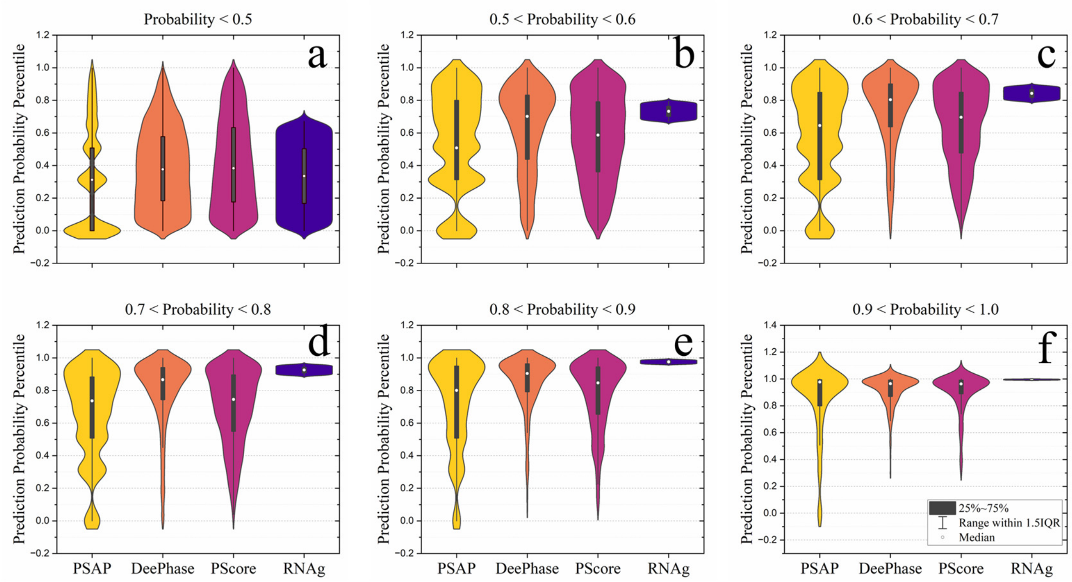


**Figure S18. Similar LLPS tendency with RNA granule tendency of identified RNA granule proteome with different prediction probabilities (0-0.5 in a, 0.5-0.6 in b, 0.6-0.7 in c, 0.7-0.8 in d, 0.8-0.9 in e and 0.9-1.0 in f) of the selected PBSG classifier.** We utilized three classic and widely applied LLPS prediction models (PSAR, DeePhase and PScore) to predict LLPS probability and applied the selected PBSG classifier to predict the RNA granule probability. Then, we calculated the average rank percentile values of prediction LLPS scores or RNA granule probability. RNAg: our RNA granule model**.**

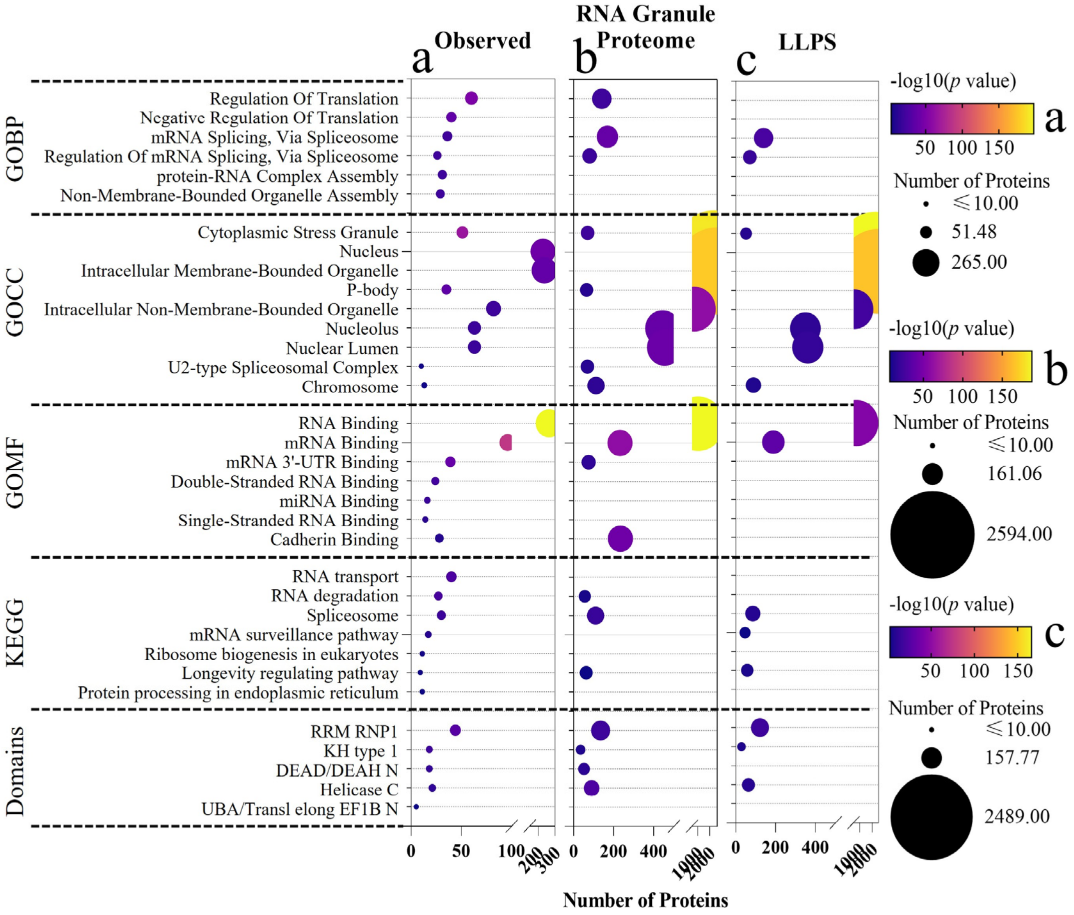


**Figure S19. Functional enrichment analyses on the observed high-confidence RNA granule proteins (a, N=429), the overall identified RNA granule proteome (b, N=6723) and the predicted LLPS-prone proteome (c, N=6723).** (a) We collected overall RNA granule proteins utilized to train our classifiers (*i.e.,* tier 1 PBSG, N=429). (b) We selected the overall RNA granule protein candidates according to the average prediction probability (over 0.5, N=6723) of the selected ten-fold evaluated PBSG classifiers. Then, we completed the functional enrichment analysis on GOBP, GOCC, GOMF, KEGG and domains of the group of proteins. (c) We utilized three classic and widely applied LLPS prediction models (PSAR, DeePhase and PScore) to select overall LLPS-prone proteins (N=6723) according to the average rank percentile values of prediction LLPS scores. The enrichment analysis is performed to compare the functional enrichment results of LLPS-prone proteins with identified and observed high-confidence RNA granule proteins. We applied the Enrichr platform to achieve the enrichment analysis.

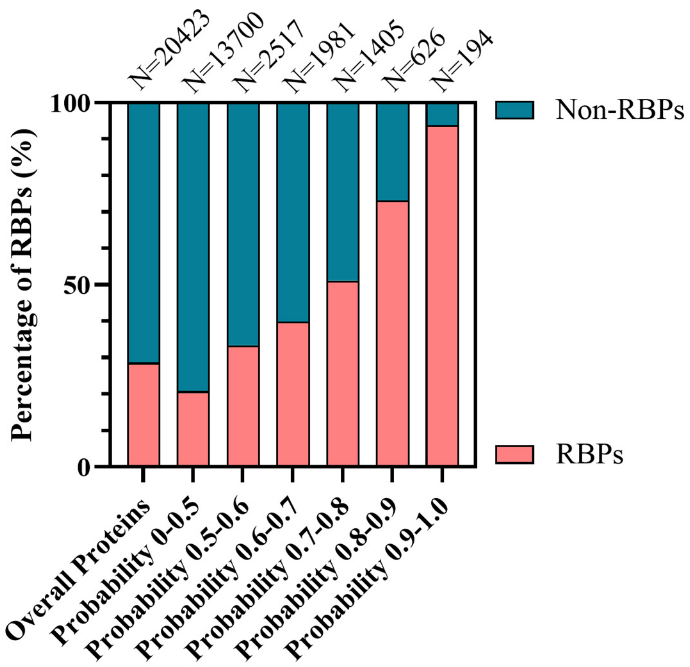


**Figure S20. Similar RBP propensities with RNA granule propensity of identified RNA granule proteome with different prediction probabilities (overall human proteome, 0-0.5, 0.5-0.6, 0.6-0.7, 0.7-0.8, 0.8-0.9 and 0.9-1.0) of the selected PBSG classifier.** We collected the human RBPs (N=6163) from the RBPbase database and applied the selected PBSG classifier to predict the RNA granule probability of each protein. Then, we calculated the percentages of RBPs in RNA granule proteome with different probabilities.

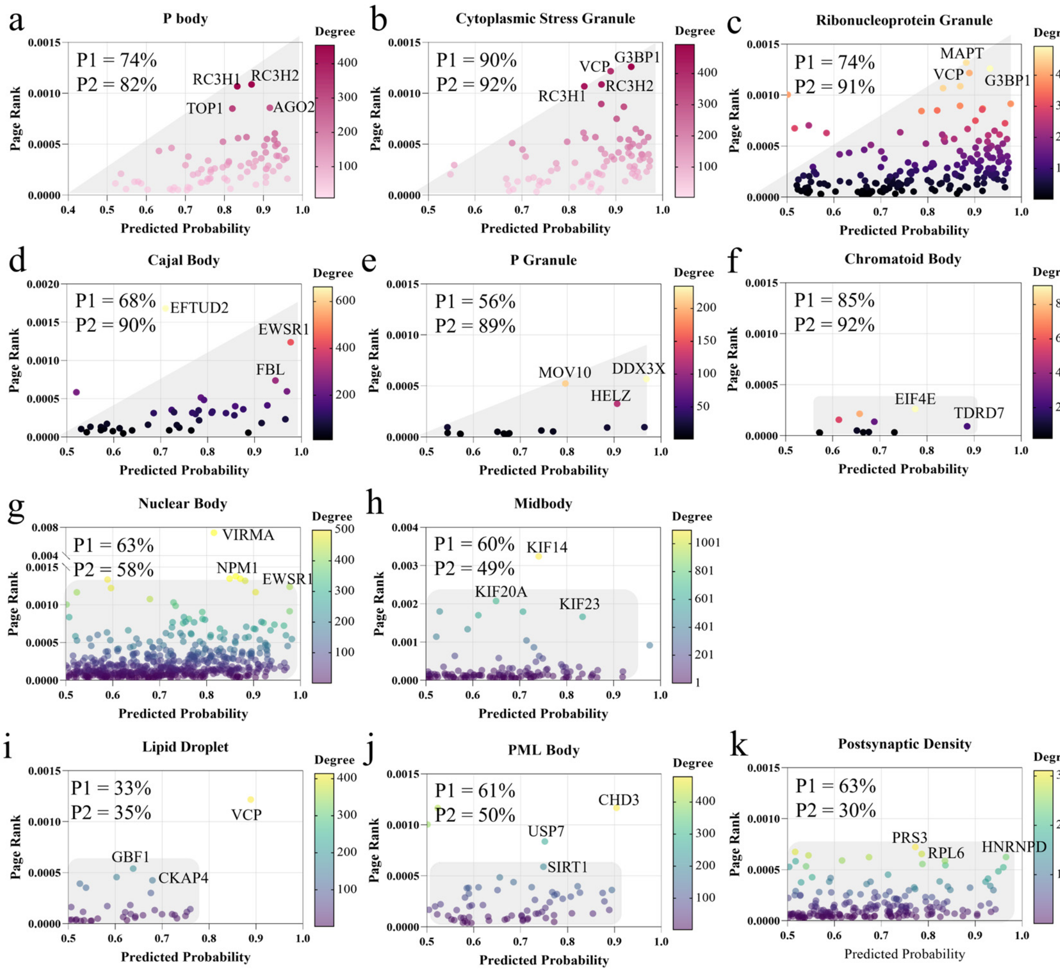


**Figure S21. Evaluation of learned component and community grammars from SG and PB on typical RNA granules and other biomolecular condensates.** We applied the predicted probability of each protein in the PBSG model to measure its likelihood to be RNA granule (*i.e.,* PB or SG) components. The PageRank value evaluates the centrality of each protein in the identified RNA granule proteome community by the PBSG model. We collected typically classified RNA granules, including PB (a), SG (b), ribonucleoprotein granule (c), Cajal body (d), P granule (e) and Chromatoid body (f), nuclear body (g), Midbody (h) and other biomolecular condensates which are not typically classified as RNA granules, including Lipid droplet (i), PML body (j) and postsynaptic density (k). We applied the distribution of protein components on the predicted probability from our PBSG model and PageRank values to evaluate the commonality of component and community grammars from PBSG models on RNA granules. P1: the percentage of protein components of each biomolecular condensate with predicted probability over 0.5 from the PBSG model. P2: the percentage of protein components of each biomolecular condensate identified as RBPs. The protein components of each biomolecular condensate were collected from the QuickGO database (Version: 2023-10-06; access date: Oct. 2023). The RBPs were collected from the RBPbase database (version: v0.2.1 alpha; access date: Oct. 2023).

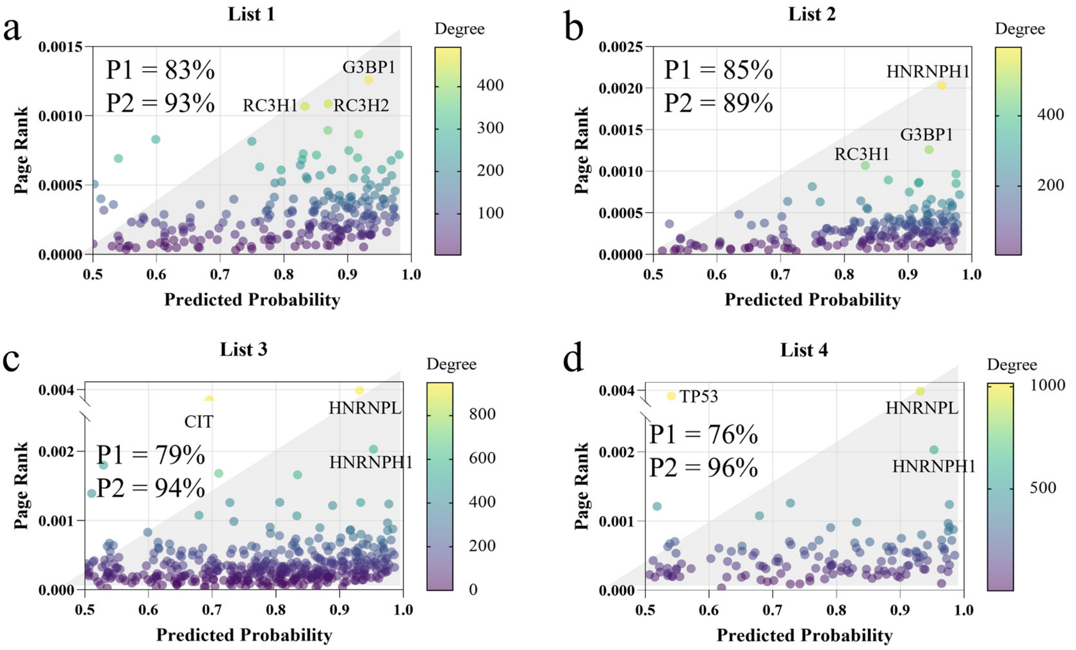


**Figure S22. The commonality evaluation of learned component and community grammars on four experimental SG proteome lists.** We applied the predicted probability of each protein in the PBSG model to measure its likelihood to be RNA granule components. The PageRank value evaluates the centrality of each protein in the identified RNA granule proteome community. We collected four experimental SG proteome lists (N = 253 in paper 1[3], N = 221 in paper 2[4], N = 486 in paper 3[5], and N = 172 in paper 4[6]. We applied the distribution of protein components on the predicted probability from our RNA granule model and PageRank values to evaluate the commonality and generalization of biomolecular and community grammars from PBSG models on experimental SG proteome. P1: the percentage of protein components of each SG proteome with predicted probability over 0.5 from the PBSG model. P2: the percentage of protein components of each SG proteome identified as RBPs. The RBPs were collected from the RBPbase database (version: v0.2.1 alpha; access date: Oct. 2023).

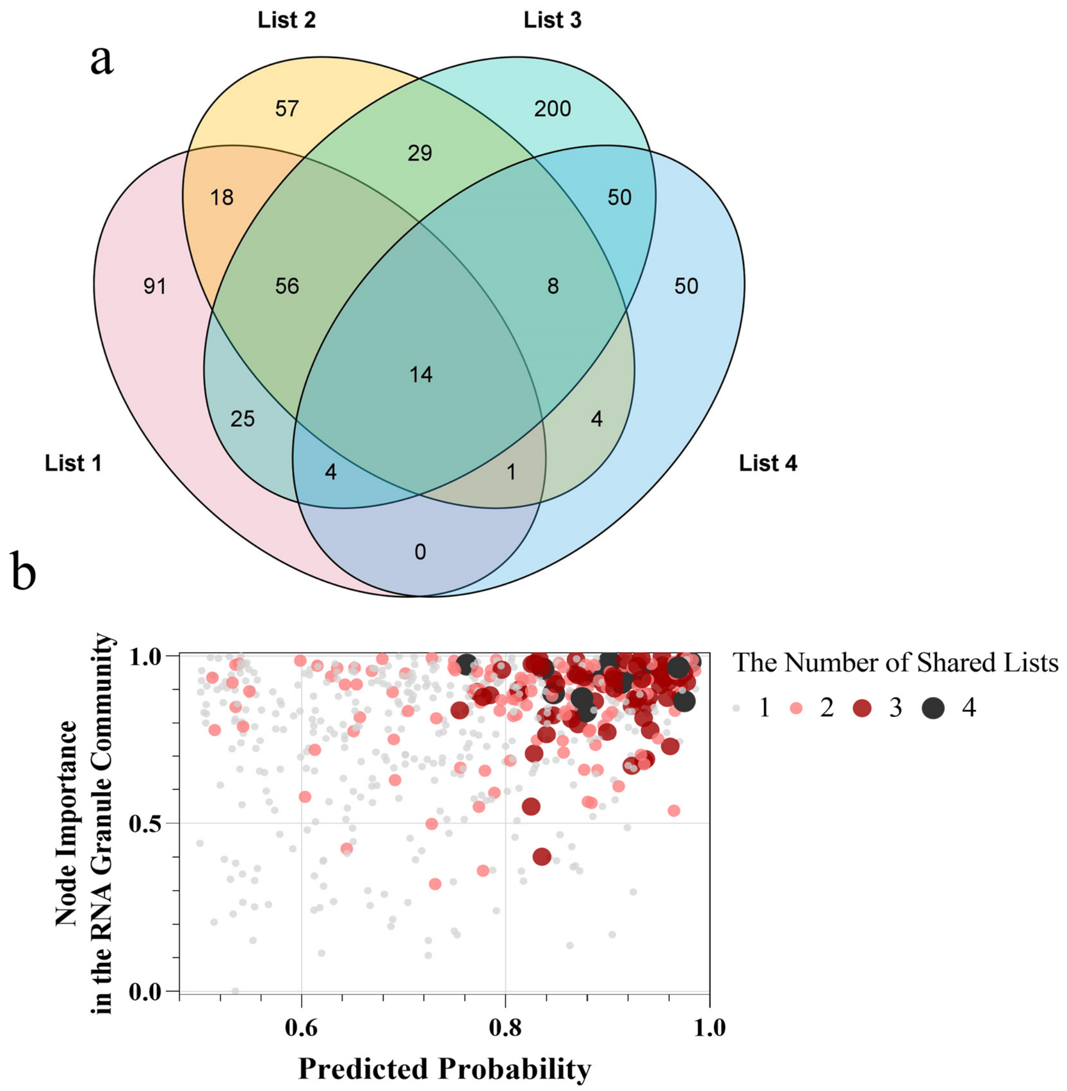


**Figure S23. Commonality evaluation of community grammars of four SG proteome collected from publications.** The Venn plot (a) illustrates the number of proteins shared in four different SG proteome. The dot plot (b) illustrates the distribution of the popular (proteins occur in most of SG proteome lists) and specific SG proteins (occurs in specific SG proteome list) on node importance and predicted probability in predicted RNA granule proteome.

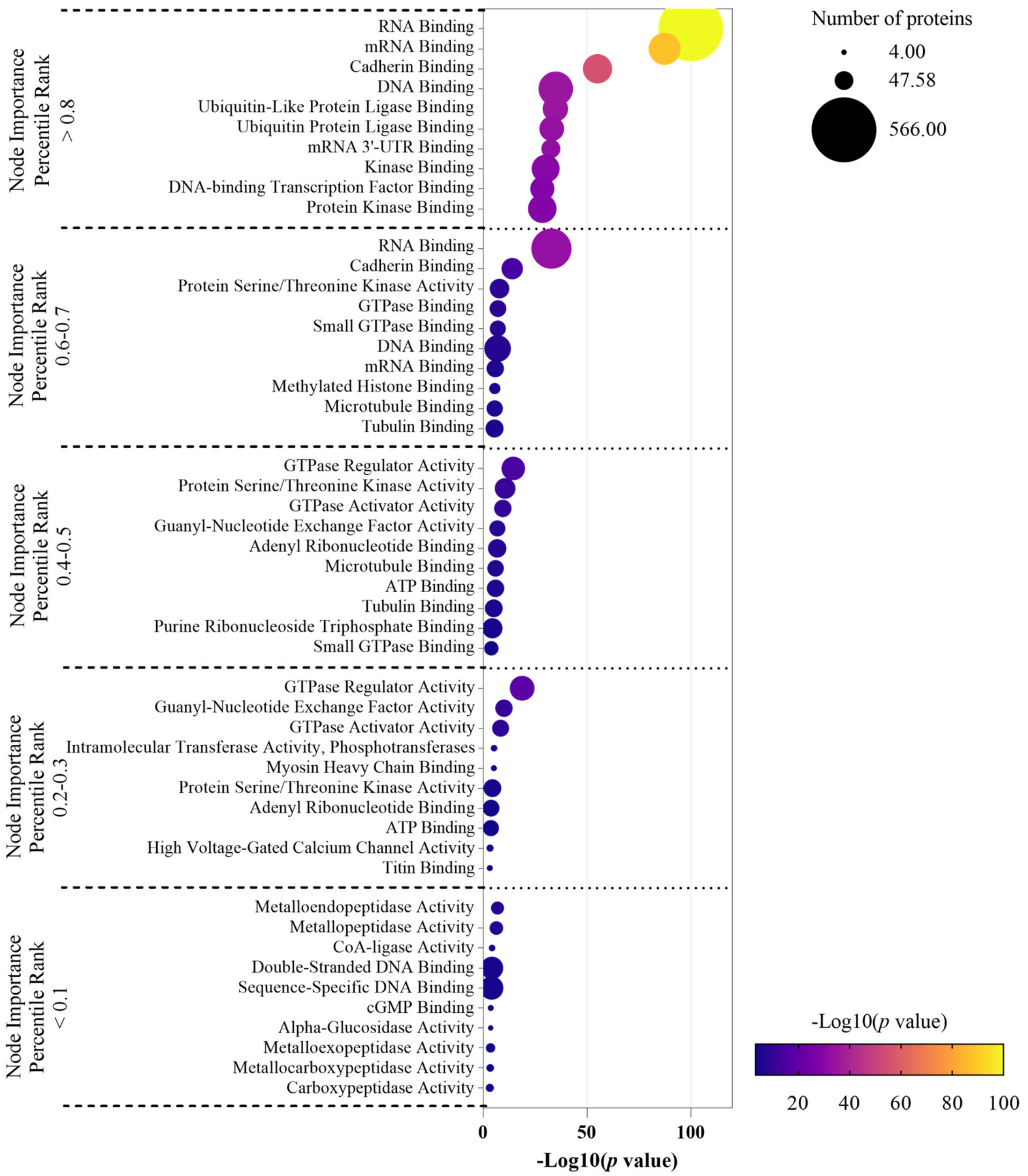


**Figure S24. The correlation between node importance in PPI network and molecular functions of predicted RNA granule proteome.** We evaluated the potential molecular functions by applying the GO enrichment molecular functions significantly ($p$ value < 0.05) enriched by proteins with different levels of node importance. The percentile rank of page rank was provided to measure importance of each protein in PPI network of the whole predicted RNA granule proteome (N=6600). We applied the Enrichr platform to achieve the enrichment analysis.

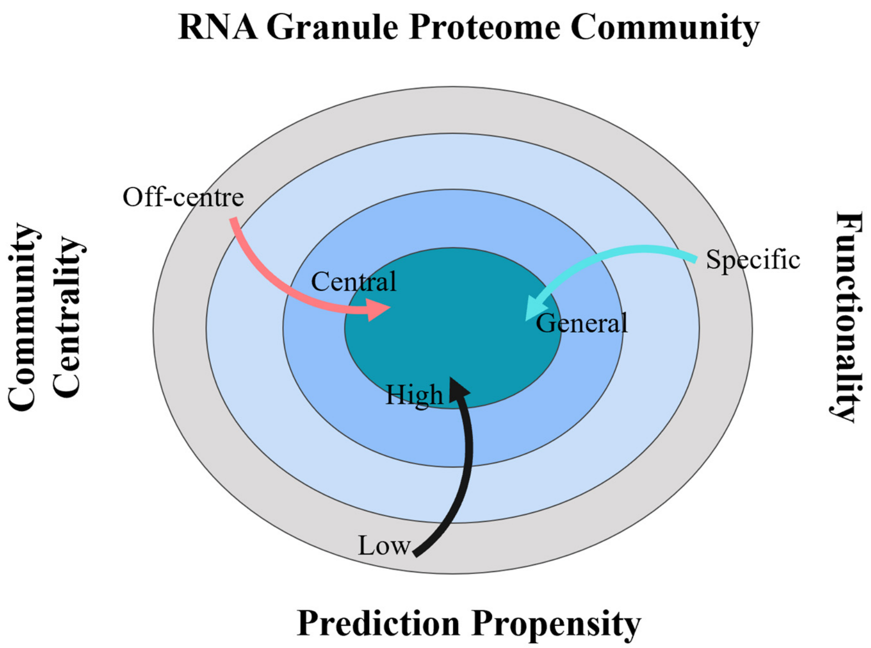


**Figure S25. Community grammars in predicted RNA granule proteome.** The community centrality represents the node importance of each protein in the RNA granule PPI community (Figure 4b-h in the main manuscript). The prediction probability represents the prediction propensity of each protein as RNA granule protein in our RNA granule model.

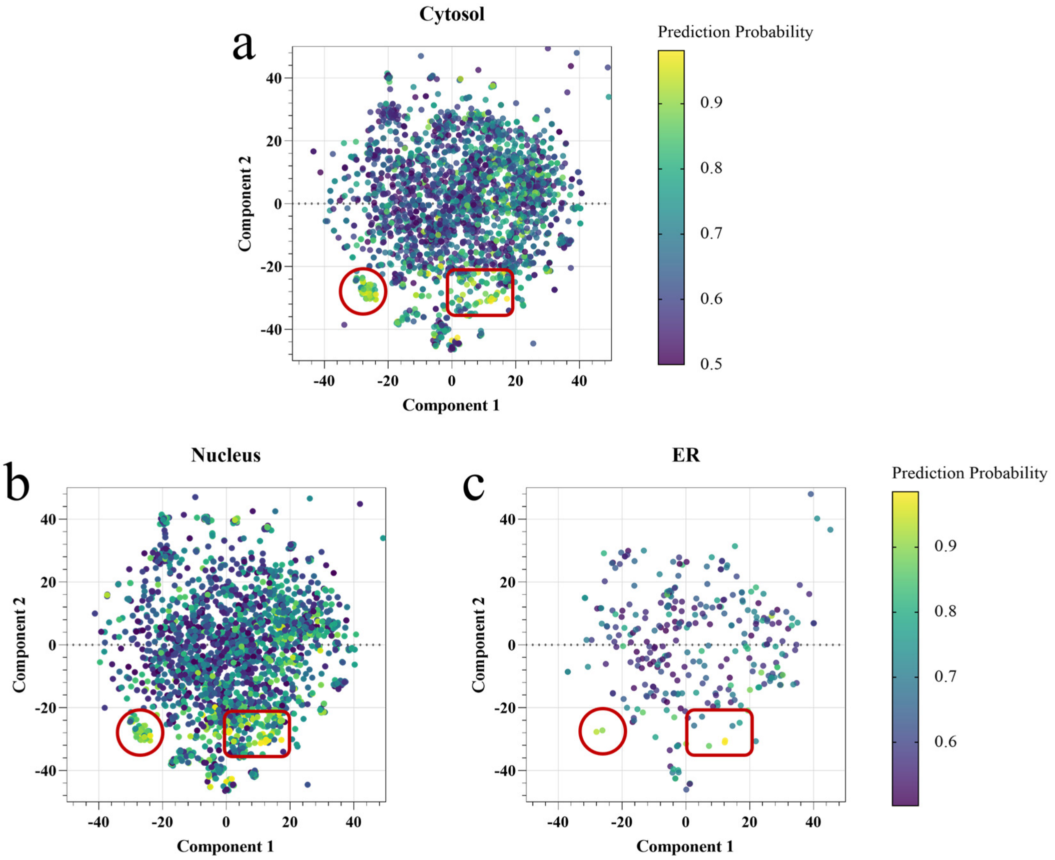


**Figure S26. Visualizations of RNA granule proteome in the cytosol, nucleus and ER by t-SNE.** We visualized overall identified RNA granule proteome PPI network (N=6600) into a 2-dimensional map and collected the locations of each protein in the overall map (*i.e.,* the component 1 and component 2 value). According to the GOquick database, we collected predicted RNA granule proteins located in cytosol (a, GO ID: 0005829, N=2652), nucleus (b, GO ID: 0005634, N=2752) and ER (c, GO ID: 0005783, N=325). We visualized the locations of protein components in cytosol (a), nucleus (b) and ER (c) using the collected protein locations of each protein in the overall map. The circle represents Cluster 1. The square represents Cluster 2. ER, endoplasmic reticulum.

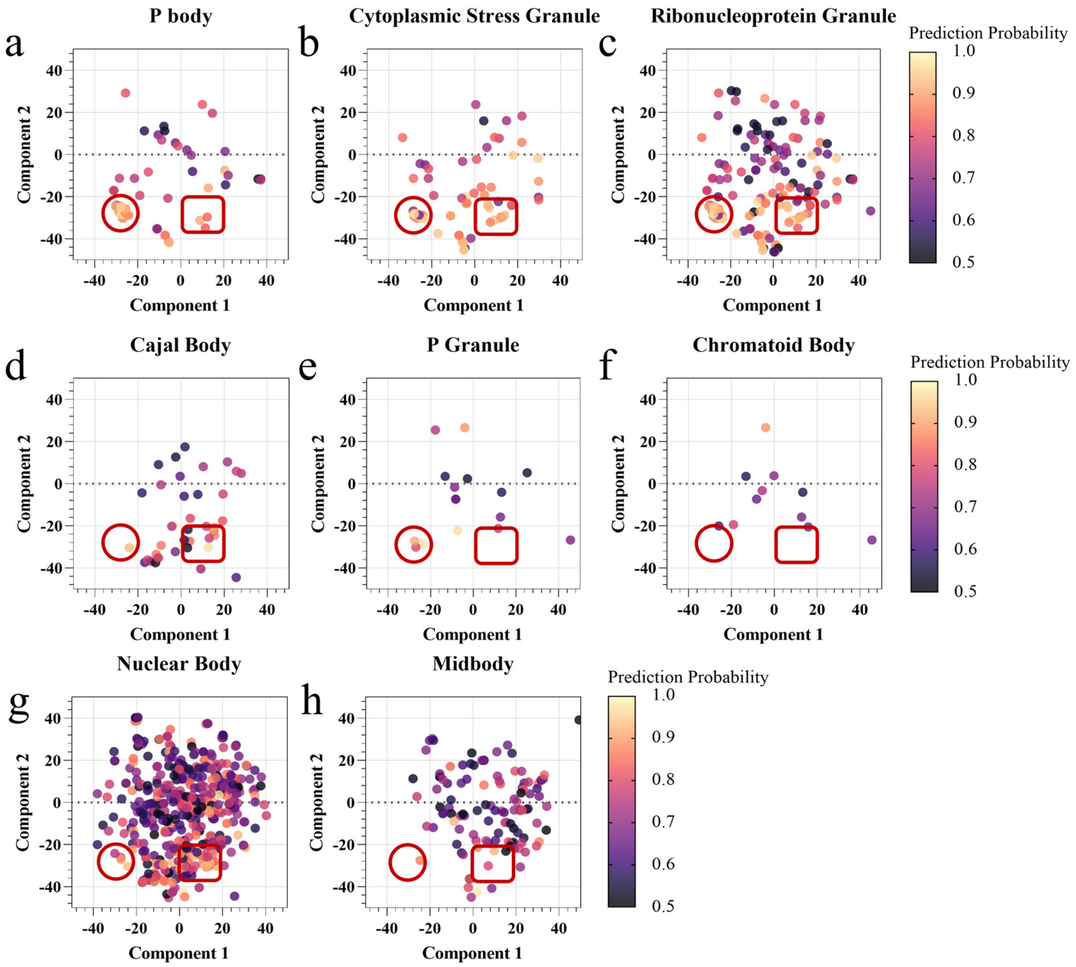


**Figure S27. Visualizations of common Cluster 1 and Cluster 2 in collected typical RNA granules by t-SNE.** (a) We visualized overall identified RNA granule proteome PPI network (N=6600) into a 2-dimensional map and collected the locations of each protein in the overall map (*i.e.,* the component 1 and component 2 value). We collected and visualized the locations of proteins of typically classified RNA granules, including PB (a), SG (b), ribonucleoprotein granule (c), Cajal body (d), P granule (e), Chromatoid body (f), nuclear body (g) and Midbody (h) using the collected protein locations in the overall map. The protein components of each typical RNA granule were collected from the QuickGO database (access date: Oct. 2023), as shown in **Error! Reference source not found.**. The circle represents Cluster 1. The square represents Cluster 2.

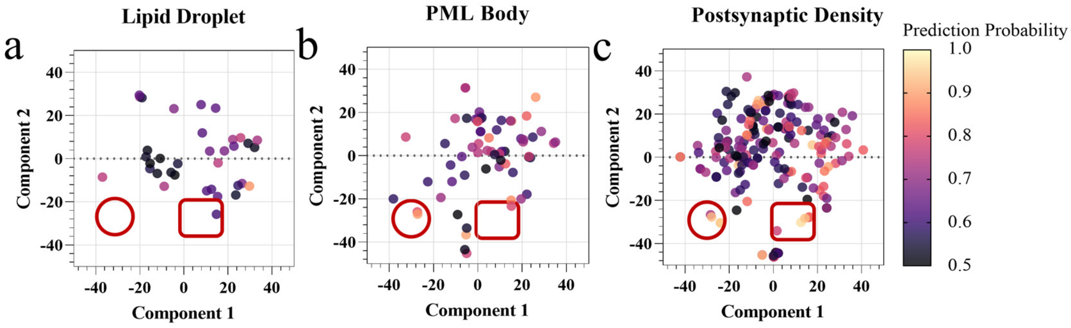


**Figure S28. Visualizations of Cluster 1 and Cluster 2 in collected non-RNA granules by t-SNE.** (a) We visualized overall identified RNA granule proteome PPI network (N=6600) into a 2-dimensional map and collected the locations of each protein in the overall map (*i.e.,* the component 1 and component 2 value). We collected and visualized the locations of proteins of typically classified non-RNA granules, including lipid droplet (a), PML body (b), postsynaptic density (c) using the collected protein locations in the overall map. The protein components of each typical non-RNA granule were collected from the QuickGO database (access date: Oct. 2023), as shown in **Table S2**. The circle represents Cluster 1. The square represents Cluster 2.

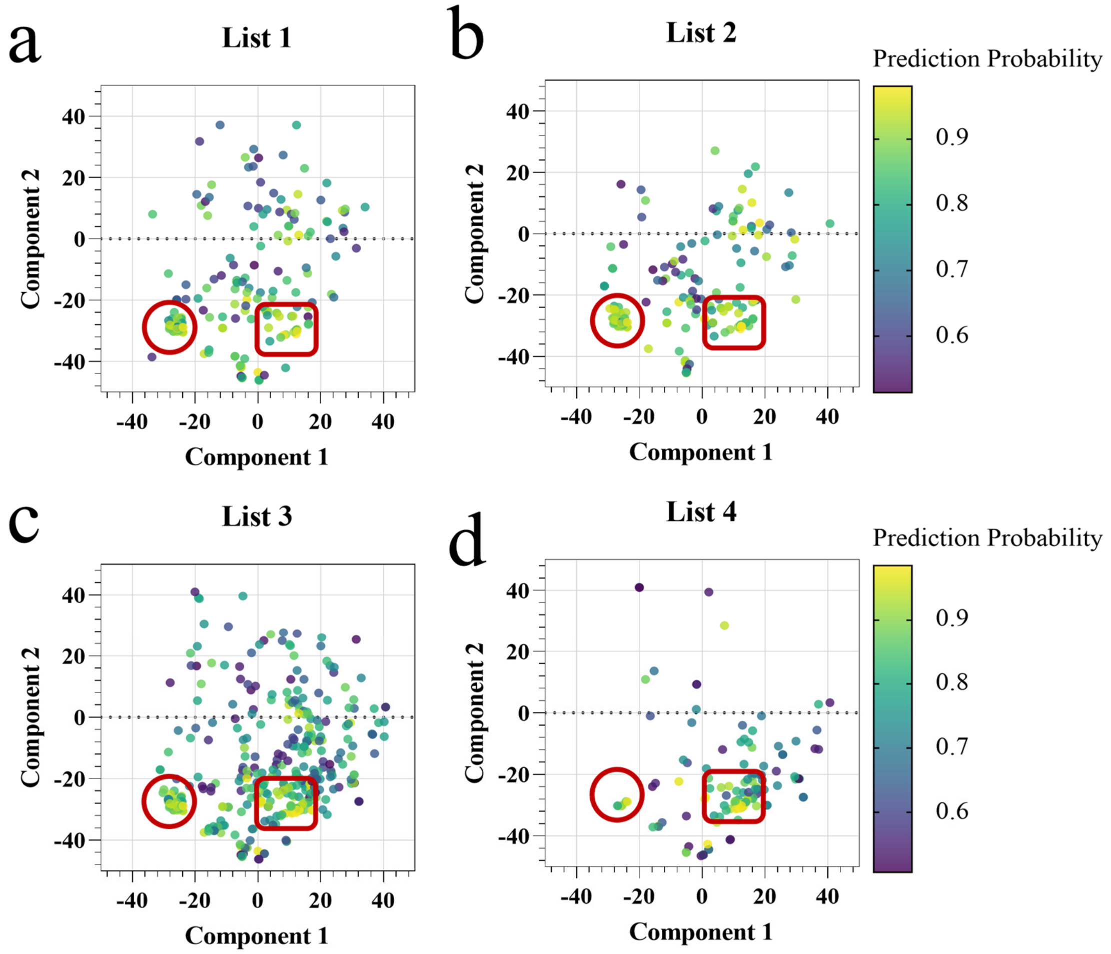


**Figure S29. Visualizations of common Cluster 1 and Cluster 2 in SG proteome lists by t-SNE.** We collected four experimental SG proteome lists (List 1: N = 253 in paper 1[3]; List 2: N = 221 in paper 2[4]; List 3: N = 486 in paper 3[5]; List 4: N = 172 in paper 4[6]. We visualized overall identified RNA granule proteome PPI network (N=6600) into a 2-dimensional map and collected the locations of each protein in the overall map (*i.e.,* the component 1 and component 2 value). We visualized the locations of protein components in List 1 (a), List 2 (b) and List 3 (c) and List 4 (d) using the collected protein locations of each protein in the overall map. The circle represents Cluster 1. The square represents Cluster 2.

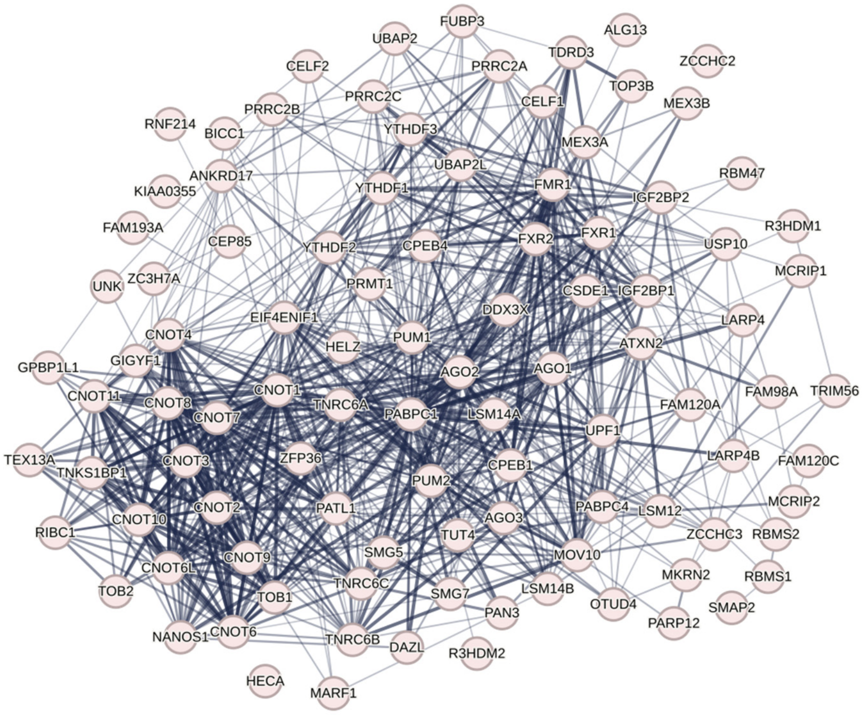


**Figure S30. Visualizations of the selected Cluster 1 subcommunity on STRING.** We collected the protein components of Cluster 1 on the overall RNA granule PPI community. Then, we visualized the PPI subcommunity of Cluster 1 proteins using STRING website. In the STRING network, the edges between two proteins represent physical and functional protein associations and the network lines between two proteins indicate the strength of data support. We built the PPI network for the extracted Cluster 1 with 100 nodes, 821 edges, 16.4 average node degree, average local clustering coefficient 0.546 and PPI enrichment $p$ value < 1.0e-16.

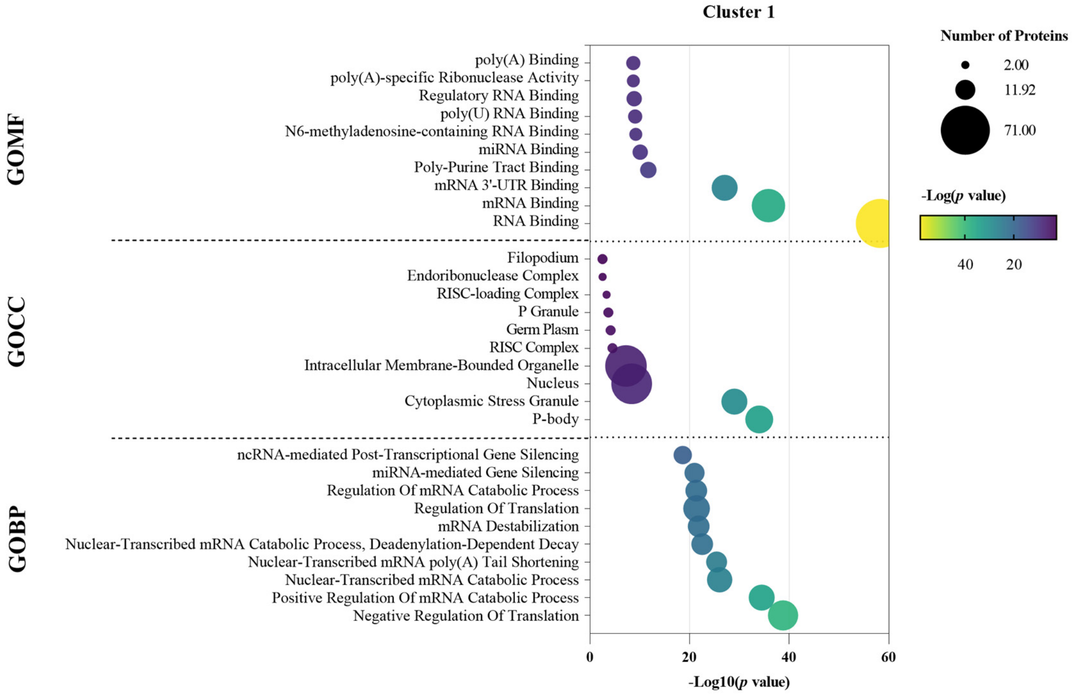


**Figure S31. The enrichment analysis of Cluster 1.** We evaluated the potential enrichment molecular functions, cellular components and biological processes by applying the GO enrichment MFs, CCs and BPs significantly ($p$ value < 0.05) enriched by extracted proteins from Cluster 1. We applied the Enrichr platform to achieve the enrichment analysis.

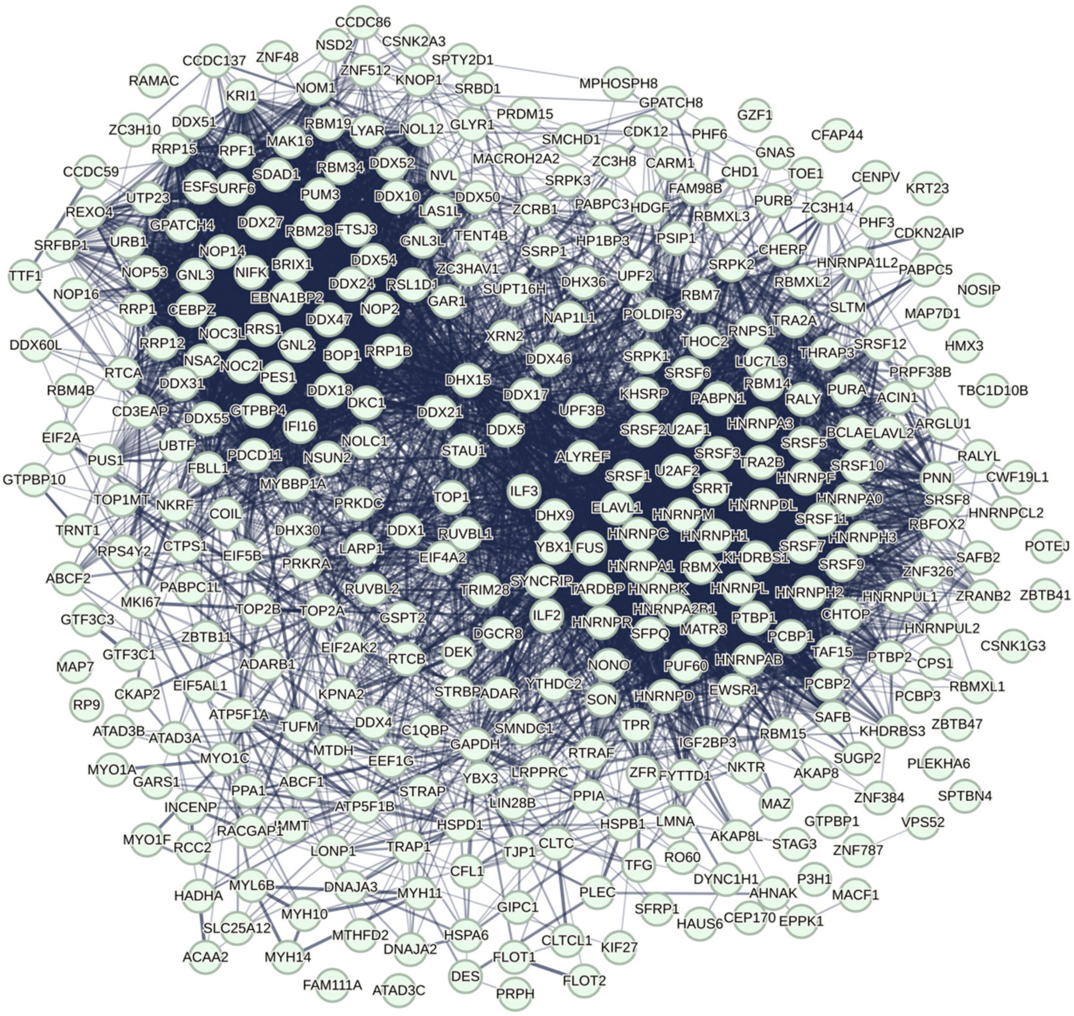


**Figure S32. Visualizations of the selected Cluster 2 subcommunity on STRING.** We collected the protein components of Cluster 2 on the overall RNA granule PPI community. Then, we visualized the PPI subcommunity of Cluster 2 proteins using STRING website. In the STRING network, the edges between two proteins represent physical and functional protein associations and the network lines between two proteins indicate the strength of data support. We built the PPI network for the extracted Cluster 2 with 351 nodes, 6163 edges, 35.1 average node degree, average local clustering coefficient 0.511 and PPI enrichment *p* value < 1.0e-16.

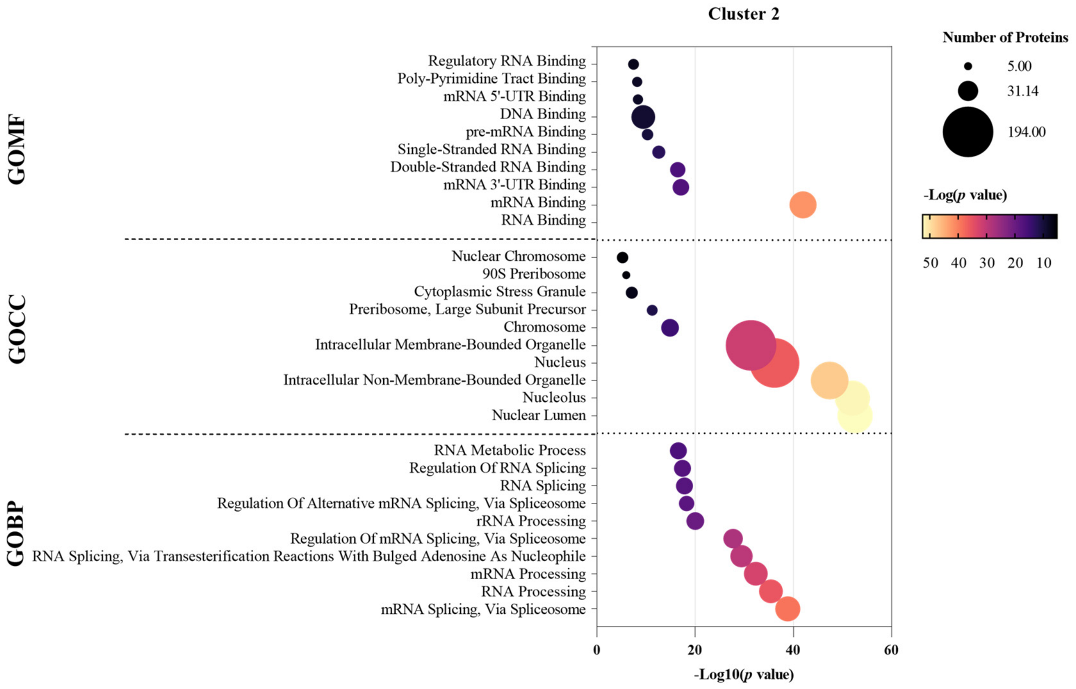


**Figure S33. The enrichment analysis of Cluster 2.** We evaluated the potential enrichment molecular functions, cellular components and biological processes by applying the GO enrichment MFs, CCs and BPs significantly ($p$ value < 0.05) enriched by extracted proteins from Cluster 2. We applied the Enrichr platform to achieve the enrichment analysis.

| **Standard** | **SG** | **PB** | **PBSG** | **$RBP_{SG}$** | **$RBP_{PB}$** | **$RBP_{PBSG}$** |
|---|---|---|---|---|---|---|
| Tier 1 | 280 | 61 | 473 | 264 | 55 | 426 |
| Tier 2 | 675 | 137 | 853 | 563 | 124 | 700 |
| Tier 3 | 399 | 189 | 854 | 288 | 82 | 501 |
| Tier 4 | - | - | 2548 | - | - | 2223 |
| Total Number | 1354 | 387 | 4728 | 1115 | 261 | 3850 |

**Table S1. Summary of human RNA granule proteins of different tiers collected from the RNAgranuleDB (access date July 2023).** $RBP_{SG}$: total number of SG proteins of each tier in the RNAgranuleDB are identified as RBPs. $RBP_{PB}$: total number of PB proteins of each tier in the RNAgranuleDB are identified as RBPs. $RBP_{PBSG}$: total number of PBSG proteins of each tier in the RNAgranuleDB are identified as RBPs.

| Prediction Probability | $N_{overall}$ | $N_{tier\_1}$ | $N_{tier\_2}$ | $N_{tier\_3}$ | $N_{tier\_4}$ | $N_{new}$ |
|---|---|---|---|---|---|---|
| 0.9-1.0 | 194 | 138 | 18 | 7 | 24 | 7 |
| 0.8-0.9 | 626 | 207 | 78 | 48 | 130 | 163 |
| 0.7-0.8 | 1374 | 108 | 159 | 103 | 334 | 670 |
| 0.6-0.7 | 1944 | 14 | 161 | 137 | 435 | 1197 |
| 0.5-0.6 | 2462 | 1 | 147 | 154 | 485 | 1675 |

**Table S2. Summary of predicted propensities of proteins in identified RNA granule PPI community by our RNA granule model and different tiers collected from the RNAgranuleDB (access date July 2023).** $N_{overall}$: total number of proteins in the prediction propensity group. $N_{tier\_1}$ to $N_{tier\_4}$: the number of the prediction propensity group in tier 1 to tier 4 proteins of the RNAgranuleDB. $N_{new}$: the number of proteins in the prediction propensity group not in the RNAgranuleDB.

| | GO ID | GO Term | Total Number of Proteins | Number of Proteins with Predicted Probability ≥ 0.5 | Percentage of Proteins with Predicted Probability ≥ 0.5 |
|---|---|---|---|---|---|
| **RNA Granules** | 0000932 | P-body | 100 | 74 | 74% |
| | 0010494 | cytoplasmic stress granule | 89 | 80 | 90% |
| | 0036464 | cytoplasmic ribonucleoprotein granule | 252 | 186 | 74% |
| | 0043186 | P granule | 27 | 15 | 56% |
| | 0033391 | chromatoid body | 13 | 11 | 85% |
| | 0015030 | Cajal body | 60 | 41 | 68% |
| | 0030496 | Midbody | 204 | 122 | 60% |
| | 0016604 | nuclear body | 833 | 521 | 63% |
| **Other Biomolecular condensates** | 0014069 | postsynaptic density | 315 | 197 | 63% |
| | 0005811 | lipid droplet | 105 | 35 | 33% |
| | 0016605 | PML body | 109 | 67 | 61% |

**Table S3. Summary of GO terms for human RNA granules and other biocondensate collected from QuickGO database (access date Oct. 2023).**

| **Extracted clusters** | $\mathbf{N_{overall}}$ | $\mathbf{N_{tier\_1}}$ | $\mathbf{N_{tier\_2}}$ | $\mathbf{N_{tier\_3}}$ | $\mathbf{N_{tier\_4}}$ | $\mathbf{N_{new}}$ |
|---|---|---|---|---|---|---|
| Cluster 1 | 100 | 91 | 2 | 1 | 0 | 6 |
| Cluster 2 | 351 | 89 | 83 | 33 | 108 | 38 |

**Table S4. Summary of extracted Cluster 1 and Cluster 2 from the identified overall RNA granule PPI community by our RNA granule model and different tiers collected from the RNAgranuleDB (access date July 2023).** $N_{overall}$: total number of proteins in the prediction propensity group. $N_{tier\_1}$ to $N_{tier\_4}$: the number of the prediction propensity group in tier 1 to tier 4 proteins of the RNAgranuleDB. $N_{new}$: the number of proteins in the prediction propensity group not in the RNAgranuleDB.
